# Ca^2+^ signaling driving pacemaker activity in submucosal interstitial cells of Cajal in the colon

**DOI:** 10.1101/2020.10.26.355404

**Authors:** Salah A. Baker, Wesley A. Leigh, Inigo F. De Yturriaga, Sean M. Ward, Caroline A. Cobine, Bernard T. Drumm, Kenton M. Sanders

## Abstract

Interstitial cells of Cajal (ICC) generate pacemaker activity responsible for phasic contractions in colonic segmentation and peristalsis. ICC along the submucosal border (ICC-SM) contributing to mixing and more complex patterns of colonic motility. We show the complex patterns of Ca^2+^ signaling in ICC-SM and the relationship between ICC-SM Ca^2+^ transients and activation of SMCs using optogenetic tools. ICC-SM displayed rhythmic firing of Ca^2+^ transients ∼15 cpm and paced adjacent SMCs. The majority of spontaneous activity occurred in regular Ca2+ transients clusters (CTCs) that propagated through the network. CTCs were organized and dependent upon Ca^2+^ entry through voltage-dependent Ca^2+^ conductances, L- and T-type Ca^2+^ channels. Removal of Ca^2+^ from the external solution abolished CTCs. Ca^2+^ release mechanisms reduced the duration and amplitude of Ca^2+^ transients but did not block CTCs. These data reveal how colonic pacemaker ICC-SM exhibit complex Ca^2+^ firing patterns and drive smooth muscle activity and overall colonic contractions.

**Synopsis:** How Ca^2+^ signaling in colonic submucosal pacemaker cells couples to smooth muscle responses is unknown. This study shows how ICC modulate colonic motility via complex Ca^2+^ signaling and defines Ca^2+^ transients’ sources using optogenetic techniques.

## Introduction

Interstitial cells of Cajal (ICC) serve several important functions in the gastrointestinal tract, including generation of pacemaker activity [1–3], neurotransduction [4–6] and responses to stretch [7]. Electrical activity in ICC are transmitted to other cells in the *tunica muscularis* via gap junctions [8]. Smooth muscle cells (SMCs), ICC and another class of platelet derived growth factor receptor alpha (PDGFRα)-positive interstitial cells are linked together in a coupled network known as the SIP syncytium [9]. The pacemaker function of ICC was deduced from morphological studies [10, 11], dissection of pacemaker regions experiments [12–14], studies on isolated ICC and SMCs [1, 15, 16], studies of muscles from animals with loss-of-function mutations in c-Kit signaling [2, 17, 18] and simultaneous impalements of ICC and SMCs [19]. While these experiments were strongly indicative of the obligatory role of ICC as pacemakers in GI smooth muscles and electrical and contractile patterns, no experiments to date have measured simultaneously pacemaker activity in ICC and responses of SMCs in terms of Ca^2+^ signaling and contraction.

The anatomy and distribution of ICC varies from place to place throughout the GI tract. Some areas have only an intramuscular type of ICC (ICC-IM) that are closely aligned with and transduce inputs from excitatory and inhibitory enteric motor neurons [4, 20–22]. Other regions contain ICC-IM and pacemaker types of ICC, that exist as a network in the myenteric plexus region of most areas of the gut (ICC-MY) [23–27]. The colon is more complex in that there are at least 4 types of ICC, distinguished by their anatomical locations and functions [28–31]. One class of colonic ICC lies along the submucosal surface of the circular muscle (CM) layer (ICC-SM). These cells are known to provide pacemaker activity in colonic muscles, and their activity is integrated with a second frequency of pacemaker activity generated by ICC-MY [12, 32–36]. Pacemaker activity generated by ICC-SM and ICC-MY causes depolarization of SMCs, generation of Ca^2+^ action potentials and excitation-contraction coupling [37].

ICC-SM generate slow waves in the canine colon [12, 38]. These are large amplitude and long duration events that produce phasic contractions [39]. The integrity of the ICC-SM network is required for regenerative propagation of slow waves, and disruption of the network causes passive decay of slow waves within a few millimeters [40]. Electrical coupling of ICC-SM into a network is an important feature allowing the pacemaker activity to coordinate the electrical activation of SMCs. ICC-SM in proximal colons of rodents also display pacemaker function, however the frequency of the slow waves is higher (10-22 min^-1^, mean 14.8. min^-1^) [35]. Slow waves, in this region of the GI tract, consist of a rapid upstroke phase, 148 mVs^-1^, that settles to a plateau phase lasting approximately 2 sec. The slow waves are coupled to low-amplitude CM contractions [37]. Colonic slow waves have been reported to depend upon both Ca^2+^ entry and intracellular Ca^2+^ release mechanisms, however Ca^2+^ signaling in colonic pacemaker cells and the coupling of Ca^2+^ events to the electrical responses were not clarified.

Previous studies have shown that all classes of ICC in the GI tract express Ca^2+^-activated Cl^-^ channels encoded by *Ano1* [41, 42]. This conductance is required for slow wave activity [42], and therefore Ca^2+^ dynamics in ICC are of fundamental importance in understanding pacemaker activity and electrical and mechanical rhythmicity in GI muscles. In the present study we tested the hypothesis that Ca^2+^ transients in ICC-SM are linked to mechanical activation of the CM and that propagation of activity in ICC-SM is related to and controlled by Ca^2+^ entry via voltage-dependent Ca^2+^ conductances. Experiments were performed on tissues containing ICC-SM taken from mice with cell-specific expression of GCaMP6f in ICC, and changes in intracellular Ca^2+^ were monitored by confocal microscopy and digital video imaging.

## Results

### ICC-SM distribution within the submucosal plexus

We optimized a preparation in which the submucosal layer was separated from the *tunica muscularis*. We found that ICC-SM were adherent to the submucosal tissues, so this preparation allowed very clear high resolution of Ca^2+^ transients in ICC-SM in the complete absence of motion artifacts due to muscle contractions. We confirmed the presence and maintenance of ICC-SM networks, which occur in intact muscles, in these preparations.

Kit immunoreactivity revealed a dense network of Kit-positive cells in the submucosal layer of the proximal colon (Fig. 1 *A*). The network consisted of ICC-SM interconnected with branching processes (Fig. 1 *B*). The average density of cell bodies was 95±28 cells mm^−2^ (*n* = 6), and the average minimum separation between cell bodies was 30.6 ± 1.8 μm (Fig. 1 *A&B*; *n* = 6). Most of ICC-SM network appeared to be adherent to the submucosal layer of the proximal colon. Isolation of the submucosa by sharp dissection showed that few ICC-SM (1.6 ± 2.1 cells mm^−2^; Fig. 1 *C*; *n* = 6) remained adherent to the muscularis in the uppermost region (1-2 μm) of the proximal colon.

**Figure 1.**
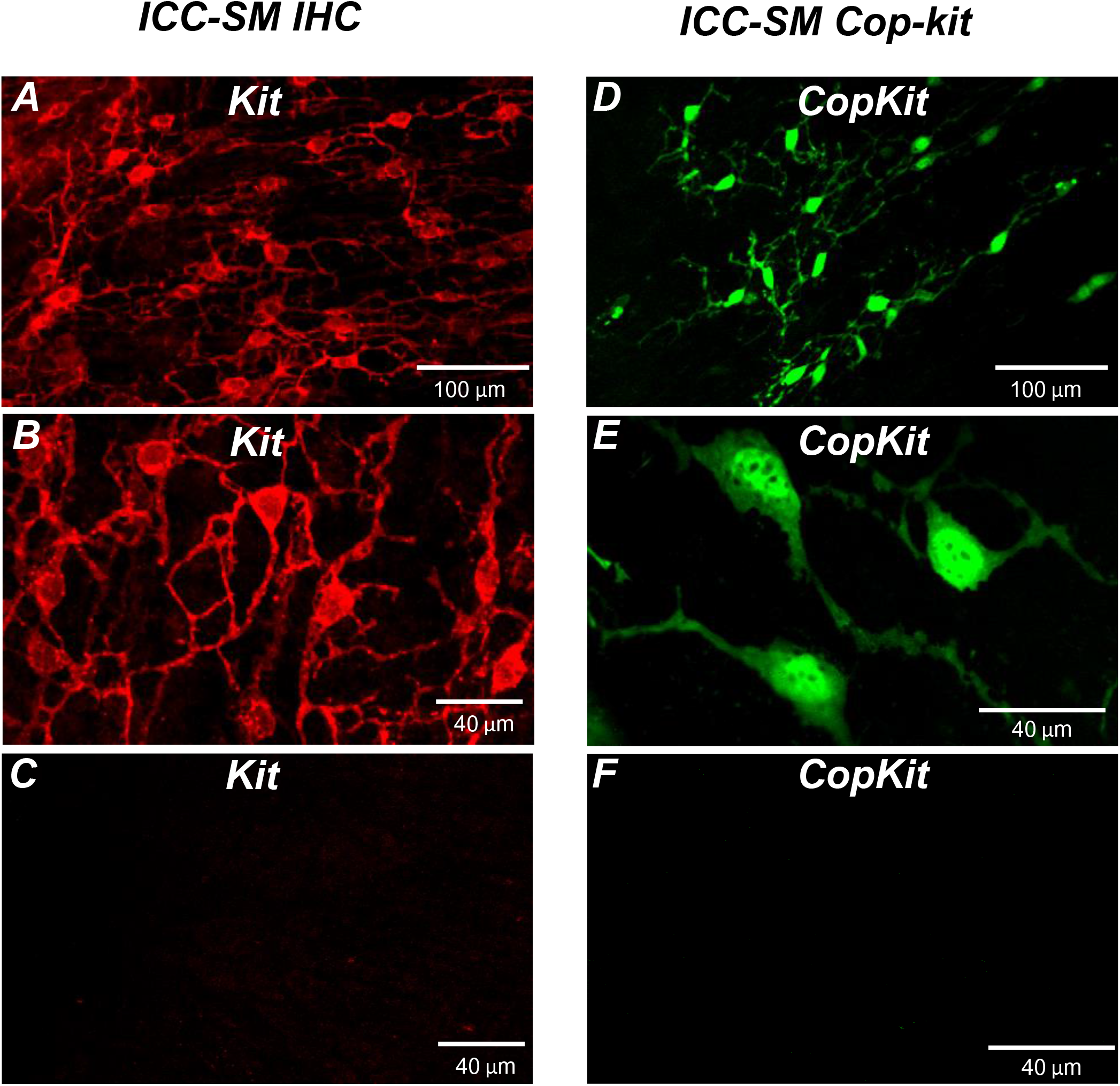
Distribution of Kit^+^ submucosal interstitial cells of Cajal (ICC-SM) in the murine colon. ***A***&***B*** Images of Kit^+^ (ICC-SM) in isolated submucosal layer of wild-type animals at ×20 and ×40 magnifications (*n* = 6). Scale bars are 100 μm and 40 μm respectively. ***C*** Absence of ICC-SM on the submucosal surface of the *tunica muscularis* of the proximal colon after removing the submucosa. ICC-SM networks are intact in preparations of submucosal tissues removed from the muscle. ***D***&***E*** ICC-SM were present in Kit^+*/copGFP*^ mice at ×20 and ×60 magnifications (*n* = 6). Scale bars are 100 μm and 40 μm respectively. ***F*** Absence of Kit^+^ (ICC-SM) on the submucosal surface of the *tunica muscularis* after removing the submucosa from Kit^+/*copGFP*^ proximal colon muscles (*n* = 6). Scale bars are 40 μm in both ***C***&***F***. All image parameters were analyzed using image J software.

Colonic muscles from Kit^+*/copGFP*^ mice expressing the copGFP exclusively in ICC were also used to confirm the distribution of ICC-SM. copGFP positive cells in the submucosal region were present at an average density of 88 ± 21 cells mm^−2^ and the average minimum separation between cell bodies was 33.7 ± 2.1 μm (Fig. 1 *D&E*; *n* = 6). ICC-SM were not resolved at the surface of the muscle layer after removing the submucosal layer (Fig. 1 *F*; *n* = 6). Colonic muscles expressing GCaMP6f exclusively in ICC were used to monitor Ca^2+^ signaling in ICC-SM. GCaMP6f positive cells in the submucosal region were present at an average density of 90 ± 32 cells mm^−2^ (*n* = 10) and the average minimum separation between cell bodies was 31.9 ± 3.2 μm. Representative images of the GCaMP6f expressing cells in ICC-SM are shown below in the Ca^2+^ imaging experiments (Fig. 2 *A*; Fig. 3 *A* and Fig. 6 *A*).

**Figure 2.**
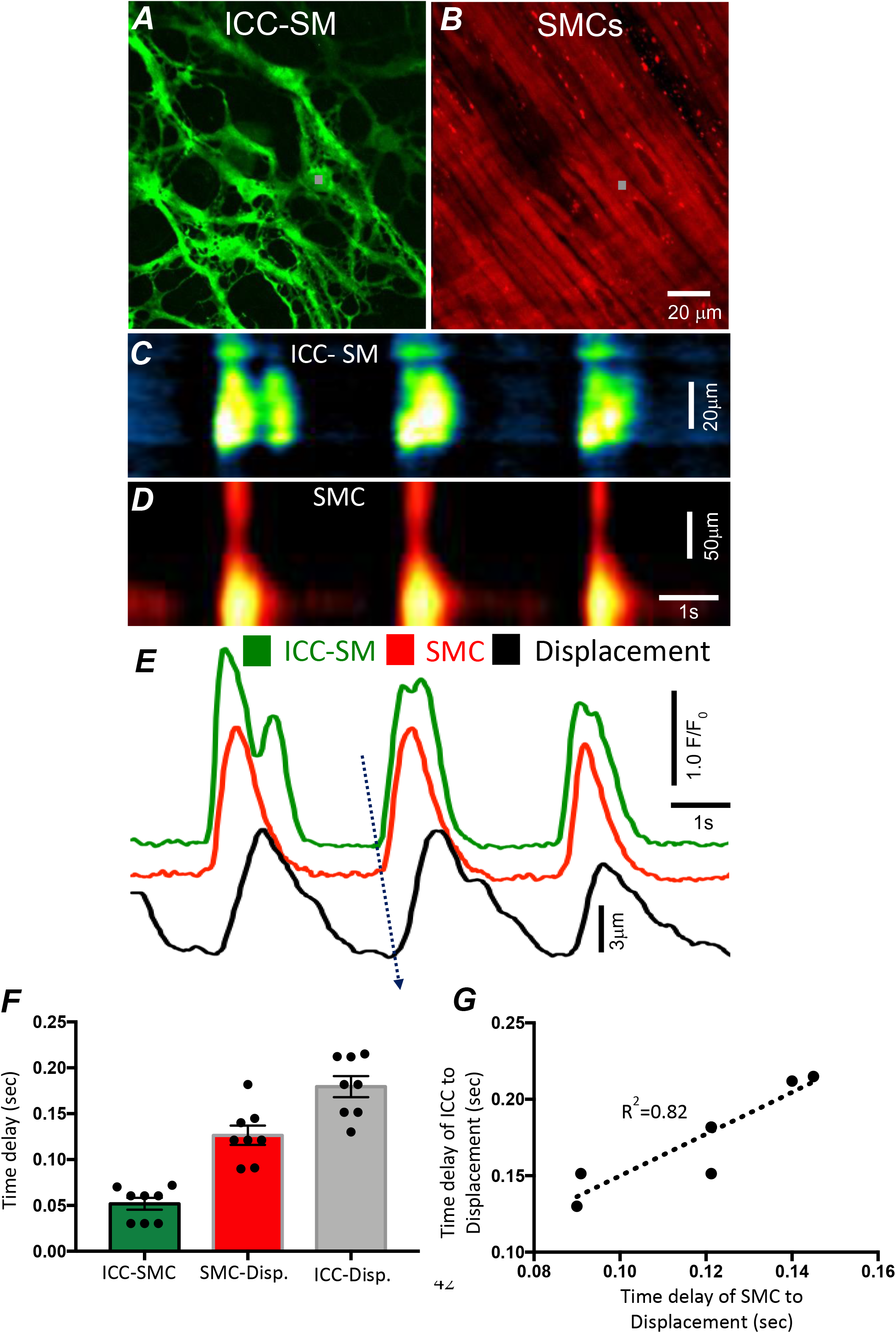
Temporal sequence of Ca^2+^ transients firing in ICC-SM and SMCs. Representative dual color imaging of ICC-SM (green; ***A***) and SMCs (red; ***B***) Ca^2+^ transients were recorded simultaneously from proximal colonic muscles of *Kit-iCre-GCaMP6f/Acta2-RCaMP1.07* mice (see Methods for details). ***C&D*** Spatiotemporal maps (STMaps) of Ca^2+^ signals in ICC-SM and SMCs during a recording period (grey boxes in ***A&B*** denote cell locations; see supplemental Movie 1). The STMaps show coordinated firing of Ca^2+^ transients in both types of cells. Ca^2+^ transient traces are plotted in ***E* (**ICC-SM-green, SMCs-red). Tissue displacement was also monitored and plotted (black trace; *E*). The black dotted arrow depicts the sequence of activation firing of Ca^2+^ transients in ICC-SM, to activation of Ca^2+^ events in SMCs, and tissue displacement. ***F*** A comparison of latencies (ms) from the start of the initiation of Ca^2+^ transients in ICC-SM to SMC activation and tissue displacement (*n* = 8). ***G*** Correlation analysis (of latencies between ICC-SM and SMC Ca^2+^ transients and tissue displacement (R^2^=0.82). All data graphed as mean ± SEM.

**Figure 3.**
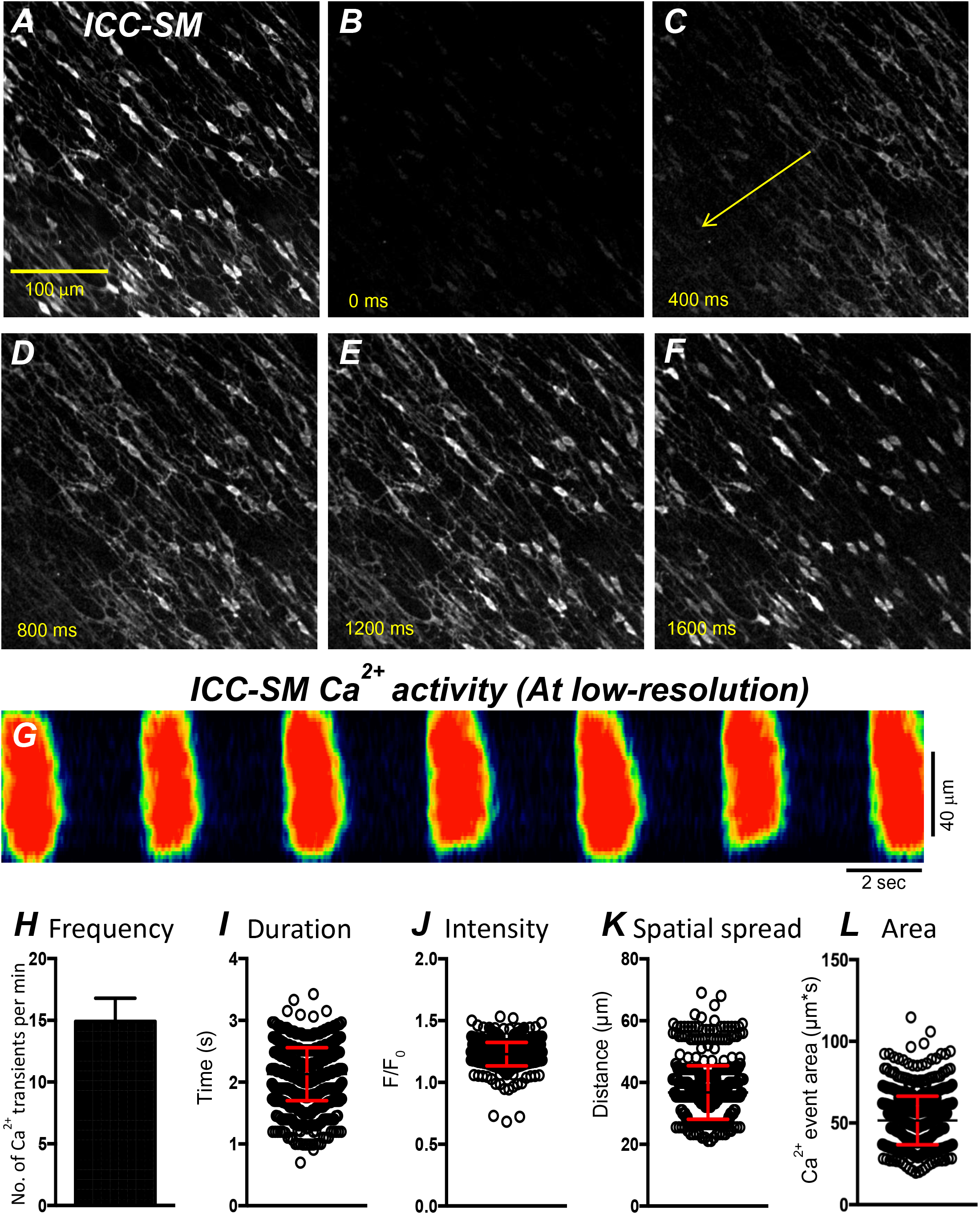
Propagating Ca^2+^ waves in ICC-SM network. ***A*** An image of ICC-SM network from the proximal colon of a Kit-iCre-GCaMP6f mouse visualized at 10x (low resolution) magnification. Scale bar is 100 μm in ***A*** and pertains to all images in ***B-F***. ***A-F*** Representative montage of the propagation of a Ca^2+^ wave throughout the ICC-SM network. The yellow arrow in panel C indicates the direction of Ca^2+^ transient propagation. ***G*** Spatiotemporal map (STMap) of Ca^2+^ signal in single ICC from the movie in panel ***A*** showing rhythmic firing of Ca^2+^ waves. Ca^2+^ activity is color coded with warm areas (red, orange) representing bright areas of Ca^2+^ fluorescence and cold colors (blue, black) representing dim areas of Ca^2+^ fluorescence. Summary data of multiple ICC-SM Ca^2+^ firing parameters (*n* = 15): frequency ***H***, duration ***I,*** intensity ***J***, spatial spread ***K*** and area of Ca^2+^ transients ***L.*** All data graphed as mean ± SEM

### Pacemaker activity of ICC-SM drives responses in SMCs

We developed a new mouse strain (*Kit-iCre-GCaMP6f/Acta2-RCaMP1.07)* that allowed simultaneous optical measurements of Ca^2+^ transients in ICC-SM and SMCs and tissue displacement (i.e. an optical means of monitoring muscle contractions). The dual color mouse allowed simultaneous imaging of the two optogenetic Ca^2+^ sensors with different fluorescence characteristics (ensuring minimal spectral overlap). The sensors were expressed in a cell-specific manner in ICC and SMCs to characterize the coordination of signaling between ICC and SMCs (Fig. 2 and supplemental movie 1). Ca^2+^ imaging of colonic muscles with attached submucosa showed a strong correlation between Ca^2+^ transients in the ICC-SM network and SMCs (Fig. 2 *A-E*; *n* = 8). Ca^2+^ transients recordings were obtained from ICC-SM and SMC at the exact coordinates in the pixel matched images (indicated by the grey box; Fig. 2 *A&B*) The sequence of activation began in ICC-SM and spread to SMCs with a latency of 56 ± 14 ms (Fig. 2 *E&F*; *n* = 8). Muscle displacement, an indicator of muscle contraction, was also measured, and displayed latencies of 120 ± 17 ms between the rise of Ca^2+^ in SMCs and resolvable displacement (Fig. 2 *E&F*; *n* = 8) and 180 ± 15 ms between activation of Ca^2+^ transients in ICC-SM and muscular contraction displacement (Fig. 2 *E&F*; *n* = 8). In few instances the time delay between ICC-SM and SMCs was not readily resolved. This could have resulted from a shift in the site of dominant pacemaker activity to a region outside the FOV. There was close correlation between the temporal latencies of ICC-SM and SMCs to tissue displacement (Fig. 2 *G*; R^2^=0.82), indicating the pacemaker nature of ICC-SM, the coupling between ICC-SM and SMCs and the management of Ca^2+^ signaling and contractions in SMCs by the pacemaker activity of ICC-SM.

### Global Ca^2+^ firing patterns in ICC-SM

Movement generated by muscle contractions is always an issue when imaging cells in muscle preparations in situ. Therefore, we prepared and utilized submucosal tissues separated from the muscle strips in most experiments. At low magnification (10x), rhythmic Ca^2+^ waves occurred and spread through ICC-SM networks in isolated submucosal layer preparations. The Ca^2+^ waves occurred at 8-22 cycle min^−1^ (Fig. 3 *A-G*) and averaged 14.9 ± 1.9 cycle min^−1^ (Fig. 3 *H*; *n* = 15; c = 120). Similar behavior occurred at similar frequencies (i.e. 16.2 ± 1.4 cycle min^−1^; *n* = 6) in ICC-SM attached to muscle tissues. Ca^2+^ waves propagated through the isolated ICC-SM networks with velocities of 219 ± 19 mm/sec. Ca^2+^ transients imaged at 10x appeared to be global and had average durations of 2.1 ± 0.4 s (Fig. 3 *I*; *n* = 15; c = 120) and amplitudes of 1.2 ± 0.3 Δ*F*/*F*_0_ (Fig. 3 *J*; *n* = 15; c = 120). The spatial spread of Ca^2+^ transients was 36.8 ± 0.4 µm (Fig. 3 *K*; *n* = 15; c = 120). The average area occupied by Ca^2+^ transients in the spatiotemporal maps was 51.6 ± 1.6 µm∗s (Fig. 3 *L*; *n* = 15; c = 120).

Cell-to-cell propagation of Ca^2+^ transients in ICC-SM is shown in Figs. 3&4. Firing of global Ca^2+^ transients appeared to be sequential in nature (Fig. 4 *A-G*; *n* = 10; c = 60), as there was a strong correlation between the occurrence of global Ca^2+^ transients in multiple ICC-SM cells (Fig. 4 *B*; *n* = 10; c = 60). Overlays of STMaps of Ca^2+^ activity in adjacent cells running parallel to each other (Fig. 4 *F&H*) showed that these Ca^2+^transients were overlapped (65.4 % of total Ca^2+^transients overlap in the FOV during a 30 s recording period) (Fig. 4 *G*). Comparison between intervals of Ca^2+^ firing of multiple ICC-SM showed strong sequential firing as each cell demonstrated very close temporal firing intervals (Fig. 4 *H&I*; *n* = 6; c = 18). These results suggest that Ca^2+^ firing in ICC-SM networks are entrained and propagate from cell-to-cell.

**Figure 4.**
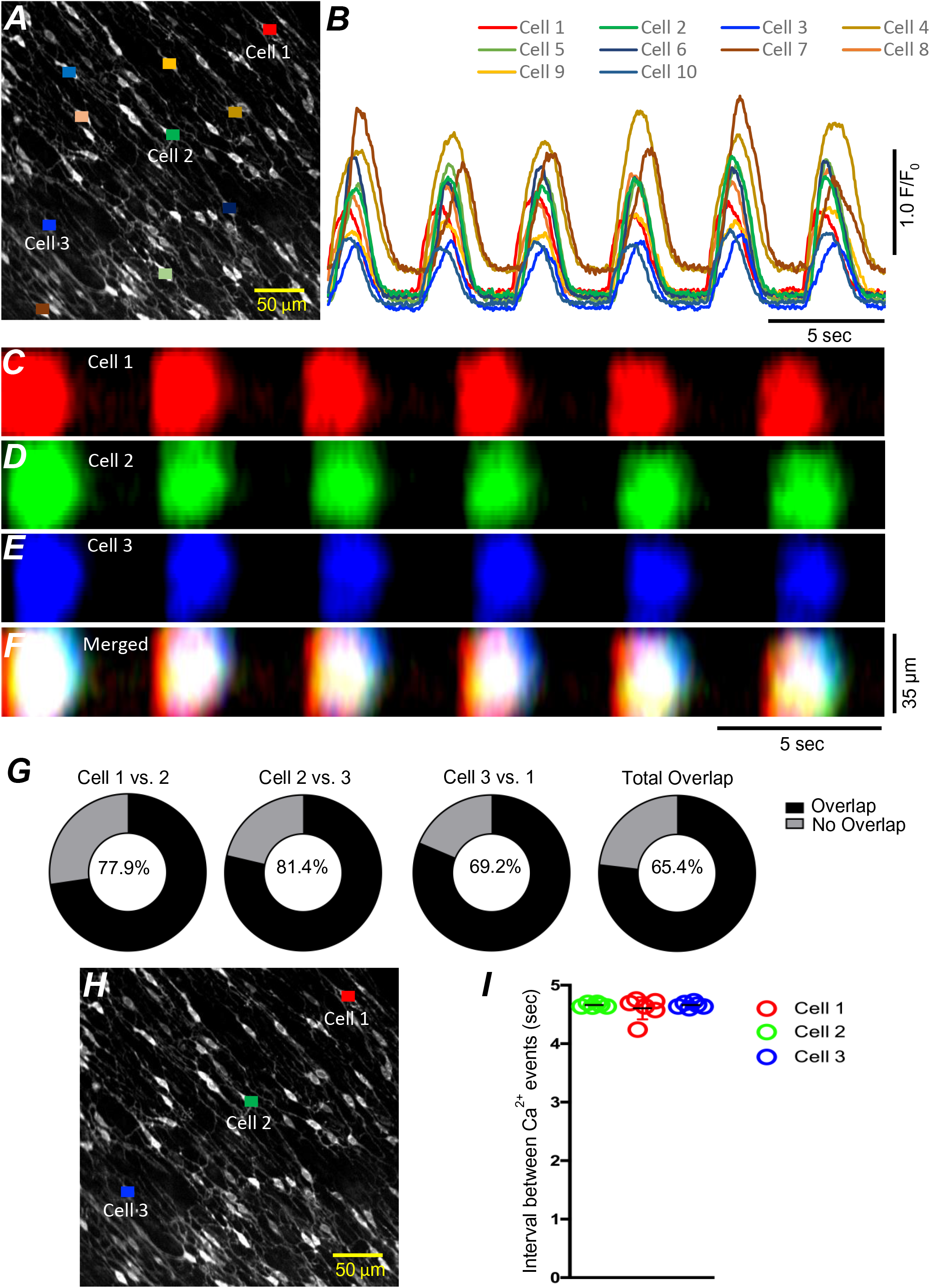
ICC-SM Ca^2+^ signaling activity is entrained. ***A*** Raw image of multiple colonic ICC-SM in a FOV. 10 cells were color-coded, and their Ca^2+^ fluorescence activity traces were plotted in ***B***. Three ICC-SM (coded as red, green and blue regions of interest; ROIs in ***A***) were selected and STMaps of the average Ca^2+^ fluorescence intensity across the diameter of the cell during a 30 s recording were constructed ***C-E***. STMaps from each cell were color coded to correspond to the red, green and blue cells and merged into a summed STMap in ***F***. Percentage of fluorescence area overlap of intracellular Ca^2+^transients between ICC-SM cells is plotted in ***G*** and cell location is identified in ***H*** (*n* = 10). The durations of Ca^2+^ waves were such that there was significant overlap of the Ca^2+^ events in individual cells across the FOVs at this magnification. Thus, each cell within the FOVs demonstrated similar temporal firing intervals (calculated as peak to peak intervals, *n* = 6) as shown in ***I*.**

### ICC-SM Ca^2+^ signals composed of multiple Ca^2+^ firing sites

Low power imaging suggested global changes in cell Ca^2+^. However, imaging of ICC-SM at higher power (60-100x) allowed more detailed visualization of the subcellular nature of Ca^2+^ transients and revealed complex firing patterns. Imaging of ICC-SM networks in situ with high spatial resolution showed that subcellular Ca^2+^ transients originated from distinct firing sites (Fig. 5; *n* = 25). STMaps constructed from Ca^2+^ transients in single ICC-SM during 30s of imaging (Fig. 5 *A&B*) identified the positions of all frequent firing sites within ICC-SM. Activity plots of individual firing sites showed they can fire once or multiple times during single Ca^2+^ waves (Fig. 5 *B&C*). The number of firing sites in a single ICC-SM ranged from 5-12 sites with an overall average of 8.2 ± 2 sites/cell (Fig. 5 *D*; *n* = 25).

**Figure 5.**
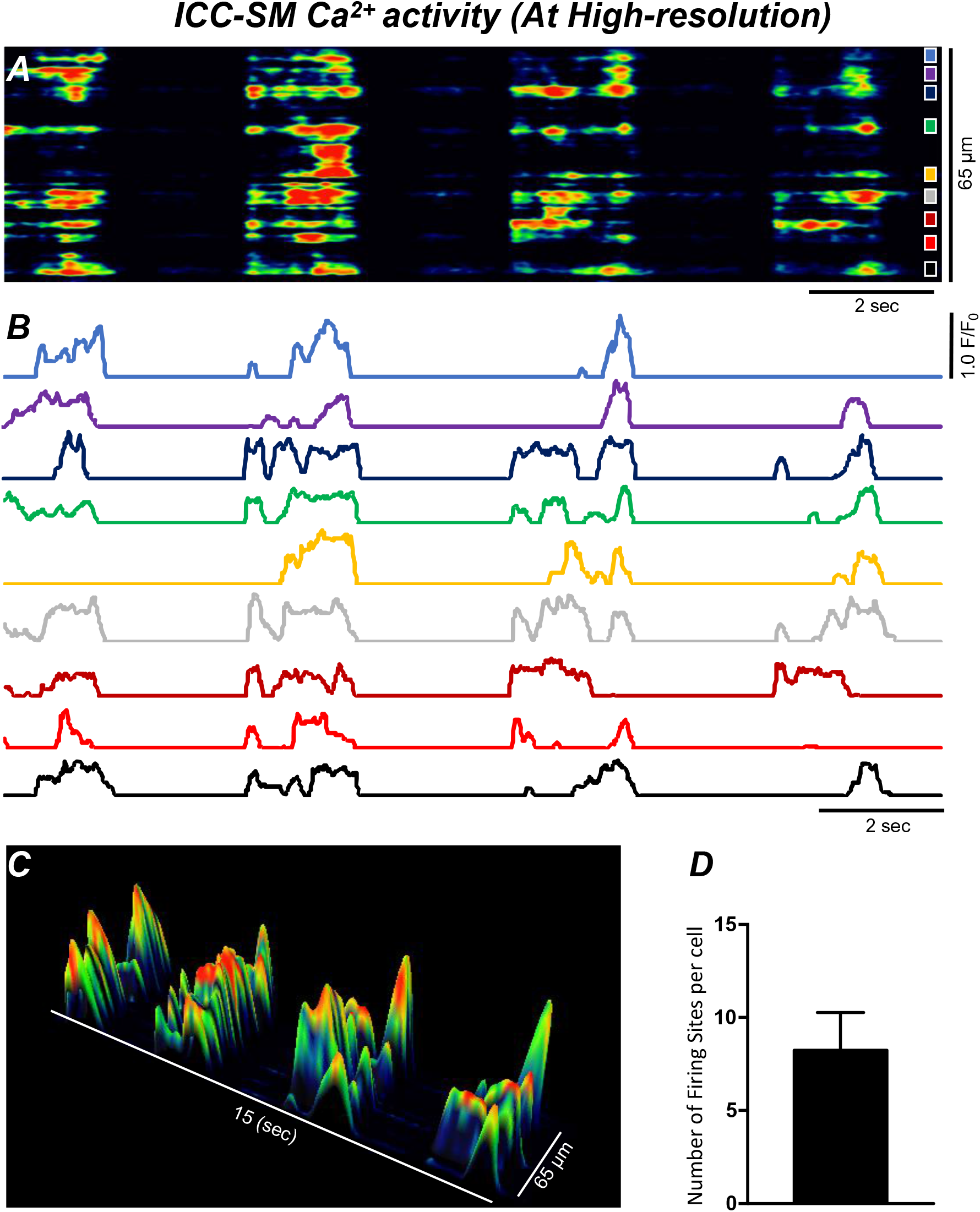
Ca^2+^ transients in ICC-SM arise from multiple firing sites. ***A*** Spatiotemporal map (STMap) of Ca^2+^ firing sites from a single ICC-SM during 4 consecutive firing cycles. Ca^2+^ activity is color coded with warm areas (red, orange) representing bright areas of Ca^2+^ fluorescence and cold colors (blue, black) representing dim areas of Ca^2+^ fluorescence. Nine distinct firing sites were detected in this cell and are marked as color squares on the right of the STMap. ***B*** Traces of the Ca^2+^ transients at each of the 9 Ca^2+^ firing sites shown on the STMap in panel ***A***. ***C*** 3-D surface plots showing the Ca^2+^ activity at the Ca^2+^ firing sites shown on the STMap in panel ***A*** over 4 consecutive Ca^2+^ firing cycles. ***D*** Average number of the firing sites per cell (*n* = 25). Data graphed as mean ± SEM.

We employed Ca^2+^ particle analysis [44] to identify and quantify Ca^2+^ firing sites in ICC-SM. Ca^2+^ firing sites were distinguished based on their spatial and temporal characteristics (Fig. 6 *A-D* and supplemental movie 2), and the sites were color-coded, shown in the example in Fig. 6 *D*, to allow visualization of active firing sites. The activities of firing sites were plotted as occurrence maps (Fig. 6 *E*). Occurrence maps of all firing sites in a network region demonstrated that global Ca^2+^ waves result from summation of localized Ca^2+^ transients from a multitude of firing sites (Fig. 6 *D-J*). The firing of Ca^2+^ transients was clustered temporally as Ca^2+^ waves swept through ICC-SM networks (Fig. 6 *E&J*). Also apparent from the occurrence maps was that the firing sequence of sites changed from Ca^2+^ wave to Ca^2+^ wave. Not all firing sites discharged Ca^2+^ transients during each wave cycle, and some sites fired more than once. From particle analysis the average particle area/frame of Ca^2+^ transients averaged 3.2 ± 0.4 µm^2^ (Fig. 6 *H*; *n* = 25) and particle count/frame averaged 0.27 ± 0.1 (Fig. 6 *I*; *n* = 25).

**Figure 6.**
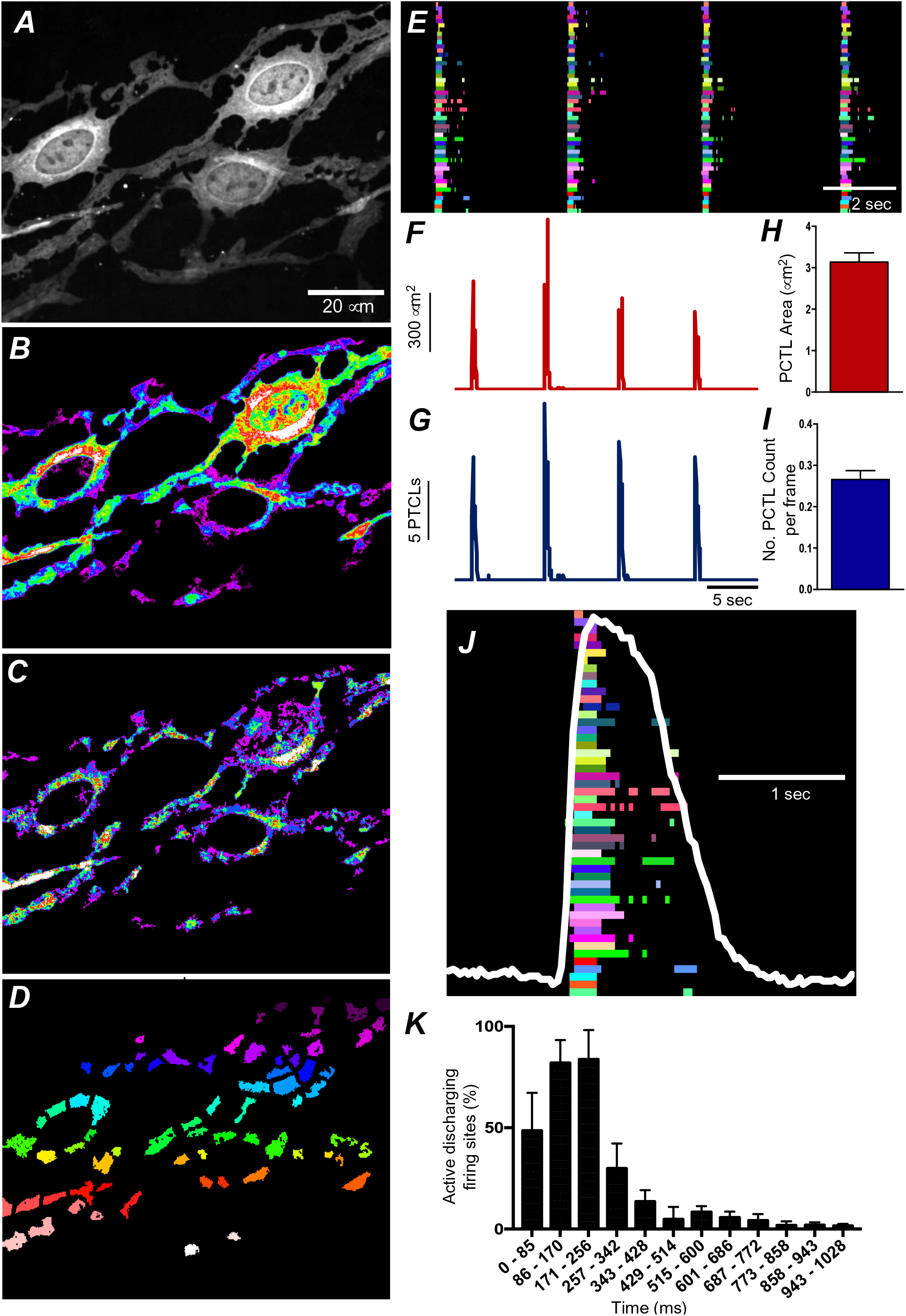
ICC-SM Ca^2+^ transient initiation sites. ***A*** Representative image of an ICC-SM network from proximal colon of a Kit-iCre-GCaMP6f mouse at 60× magnification. ***B*** Heat map of total Ca^2+^ PTCLs generated from the video shown in ***A*** (see Supplemental Movie 2). ***C*** Particles were threshholded temporally to generate a heat map indicating Ca^2+^ firing sites in the ICC-SM. ***D*** Image showing individually color-coded Ca^2+^ firing sites in the FOV shown in ***C***. ***E*** The temporal characteristics of each individual, color-coded firing site is displayed as an occurrence map, with each “lane” representing the occurrence of firing PTCLs within each firing site. Activity traces for PTCLs for the duration of recording from the entire FOV are shown in ***F&G***. Traces for PTCL area (***F***; red) and PTCL count (***G***; blue) are shown. ***H&I*** Summary graphs show averag***e*** PTCL areas and counts for Ca^2+^ firing sites in ICC-SM (*n* = 25). ***J*** One Ca^2+^ wave in ICC-SM (white trace) is expanded and the numerous Ca^2+^ initiation sites that fired during this wave are superimposed. ***K*** Distribution plot showing average percentages of firing sites during a Ca^2+^ wave. Values are calculated for 1s and plotted in 85 ms bins (*n* = 25). Data graphed as mean ± SEM.

Particle analysis also showed that Ca^2+^ firing sites were most active during the first ∼256 ms of a Ca^2+^ wave, and activity decayed with time. This point was further illustrated by distribution plots showing the average percentage of firing sites discharging at various times during Ca^2+^ waves (Fig. 6 *K*; *n* = 25). Initially high firing and decay as a function of time suggest that Ca^2+^ entry mechanisms may be important for: i) initiating Ca^2+^ transients, and ii) organizing the occurrence of Ca^2+^ transients into clusters. It is also possible that Ca^2+^ stores are loaded during the diastolic period between Ca^2+^ waves and therefore more excitable at the onset of a Ca^2+^ wave.

### Ca^2+^ influx mechanisms are required for initiation of clustered Ca^2+^ transients in ICC-SM

The effects of reducing extracellular Ca^2+^ ([Ca^2+^]_o_) on Ca^2+^ transients was investigated to evaluate the importance of Ca^2+^ influx for the patterning of Ca^2+^ signaling in ICC-SM. Removal of [Ca^2+^]_o_ abolished Ca^2+^ transients in ICC-SM within 10 min (Fig. 7 *E-L*; *n* = 6). Stepwise reduction in [Ca^2+^]_o_ (from 2.5 mM to 0 mM) showed that Ca^2+^ transients decreased in a concentration-dependent manner (Fig. 7*; n* = 6). ICC-SM Ca^2+^ PTCL area and count of firing sites were reduced in response to lowering [Ca^2+^] _o_ (Fig. 7 *F-J*; *n* = 6). Reducing [Ca^2+^]_o_ from 2.5 mM (control conditions) to 2 mM caused a reduction in the average PTCL area by 19.6 ± 4.5 % (Fig. 7 *K*; *n* = 6) and PTCL count by 25.4 ± 2.9 % (Fig. 7 *L*; *n* = 6). Further reduction of [Ca^2+^] _o_ to 1 mM reduced Ca^2+^ transient parameters by 47.2 ± 3.5 % PTCL area (Fig. 7 *K*; *n* = 6) and 45.1 ± 4.4 % PTCL count (Fig. 7 *L*; *n* = 6). Lowering [Ca^2+^] _o_ to 0.5 mM also reduced Ca^2+^ PTCL area by 73.9 ± 3 % (Fig. 7 *K*; *n* = 6) and PTCL count by 73.2 ± 2 % (Fig. 7 *L*; *n* = 6). Removal of Ca^2+^ from the extracellular solution (Ca^2+^-free KRB solution containing 0.5 mM EGTA) abolished Ca^2+^ signals 8-10 min after solution replacement. The Ca^2+^ transient PTCL area was reduced by 99.4 ± 0.6 % (Fig. 7 *K*; *n* = 6) and PTCL count by 99.3 ± 0.7 % (Fig. 7 *L*; *n* = 6). We also noted that the highly organized CTCs occurring in the presence of 2.5 mM [Ca^2+^] _o_ became less organized as [Ca^2+^] _o_ was reduced (e.g. compare the tight clusters in Fig. 7A with the more diffuse clusters in Fig. 7C). These experiments highlight the importance of Ca^2+^ entry mechanisms in Ca^2+^ signaling within ICC-SM.

**Figure 7.**
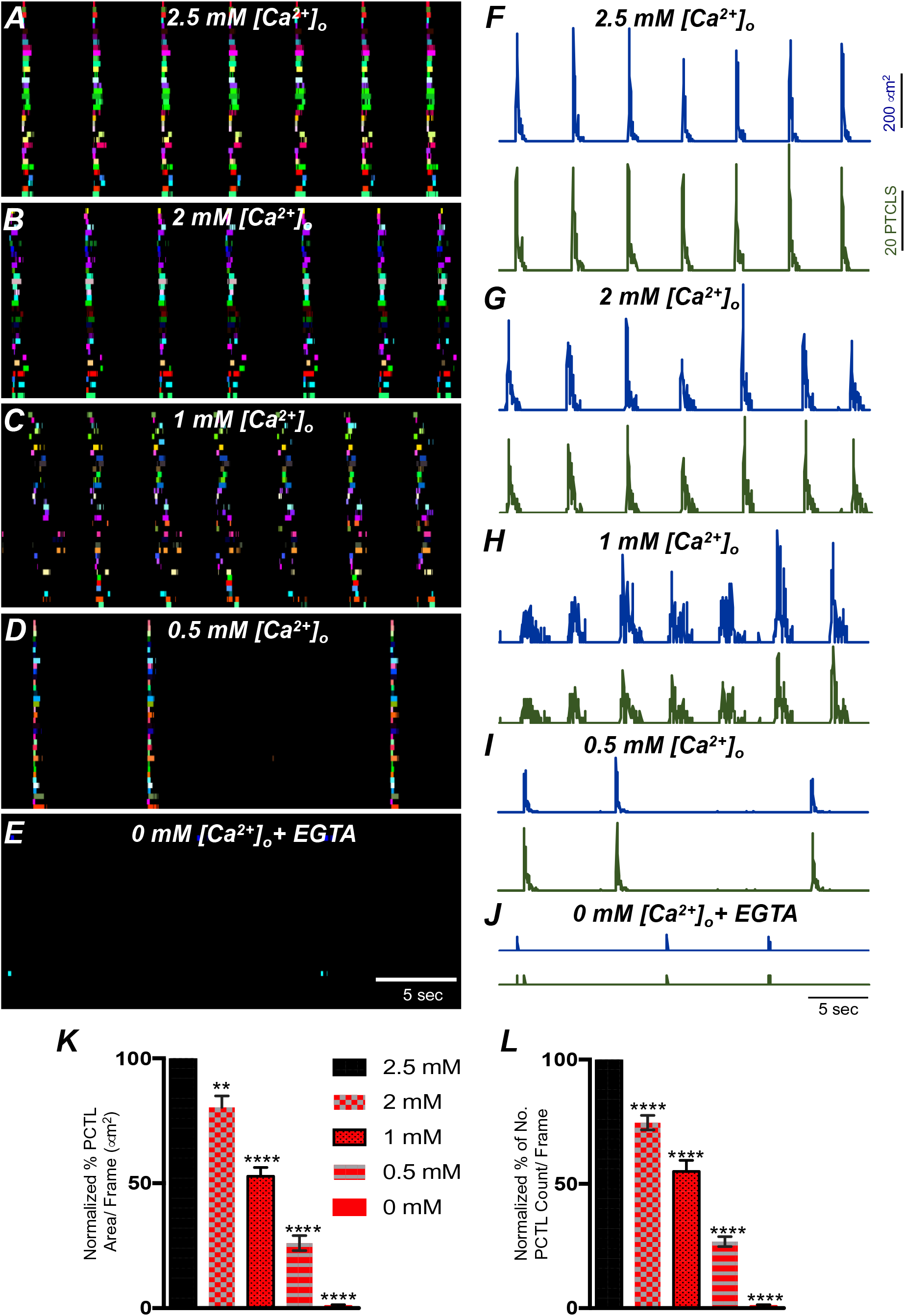
Effects of [Ca^2+^]_o_ Ca^2+^ transients in ICC-SM. ***A*** ICC-SM Ca^2+^ transients and Ca^2+^ firing sites were color-coded and plotted as an occurrence map under control conditions with [Ca^2+^]_o_ = 2.5 mM. ***B-E*** showing the effects of reducing [Ca^2+^]_o_ to 2 mM ***B***; 1 mM ***C***; 0.5 mM ***D*** and after Ca^2+^ removal of [Ca^2+^]_o_ (0 mM added and final solution buffered with 0.5 mM EGTA) ***E***. ***F-J*** Traces of Ca^2+^ PTCL activity in ICC-SM (PTCL area, blue and PTCL count, green) under control conditions ***F*** and after reducing [Ca^2+^]_o_ to 2 mM ***G***; 1 mM ***H***; 0.5 mM ***I*** and after removal of [Ca^2+^]_o_ as shown in ***J***. Summary graphs of Ca^2+^ PTCLs in ICC-SM under control conditions and with reduced [Ca^2+^]_o_ in ***K*** (PTCL area) and ***L*** (PTCL count). Data were normalized to controls and expressed as percentages (%). Significance was determined using one-way ANOVA, ** = *P*<0.01, **** = *P*<0.0001, *n*=6. All data graphed as mean ± SEM.

### Molecular expression of Ca^2+^entry channels in ICC-SM

The apparent dependence on the Ca^2+^ gradient to maintain pacemaker function in ICC-SM suggests that Ca^2+^ entry mechanisms are critical for initiation and organization of CTCs. Therefore, we examined expression of several Ca^2+^ channels that might convey Ca^2+^ entry in ICC-SM (Fig. 8). After isolation of the submucosal layer from the proximal colon of Kit^+/copGFP^ mice and subsequent cell dispersion, we sorted copGFP-positive ICC-SM with fluorescence activated cell-sorting (FACS) and evaluated the expression of voltage-dependent and-independent Ca^2+^ channels by qPCR (Fig. 8 *A&B*). First, we confirmed the purity of sorted ICC-SM with cell-specific markers (Fig. 8 *A*). Kit receptors and ANO1 channels are signatures of ICC throughout the GI tract and enrichment of *Kit* and *Ano1* expression was observed in sorted ICC-SM compared to unsorted cells (*Kit* expression was 0.21 ± 0.014; *Ano1* expression was 0.14 ± 0.02 relative to *Gapdh*). The expression levels of *Myh11* a smooth muscle cell marker and *Uch11* a panneuronal marker encoding PGP9.5 were minimal (*Myh11* expression was 0.03 ± 0.002; Uch*11* expression was 0.002 ± 0.0001 relative to *Gapdh*). confirming the purity of ICC-SM sorted by FACS.

**Figure 8.**
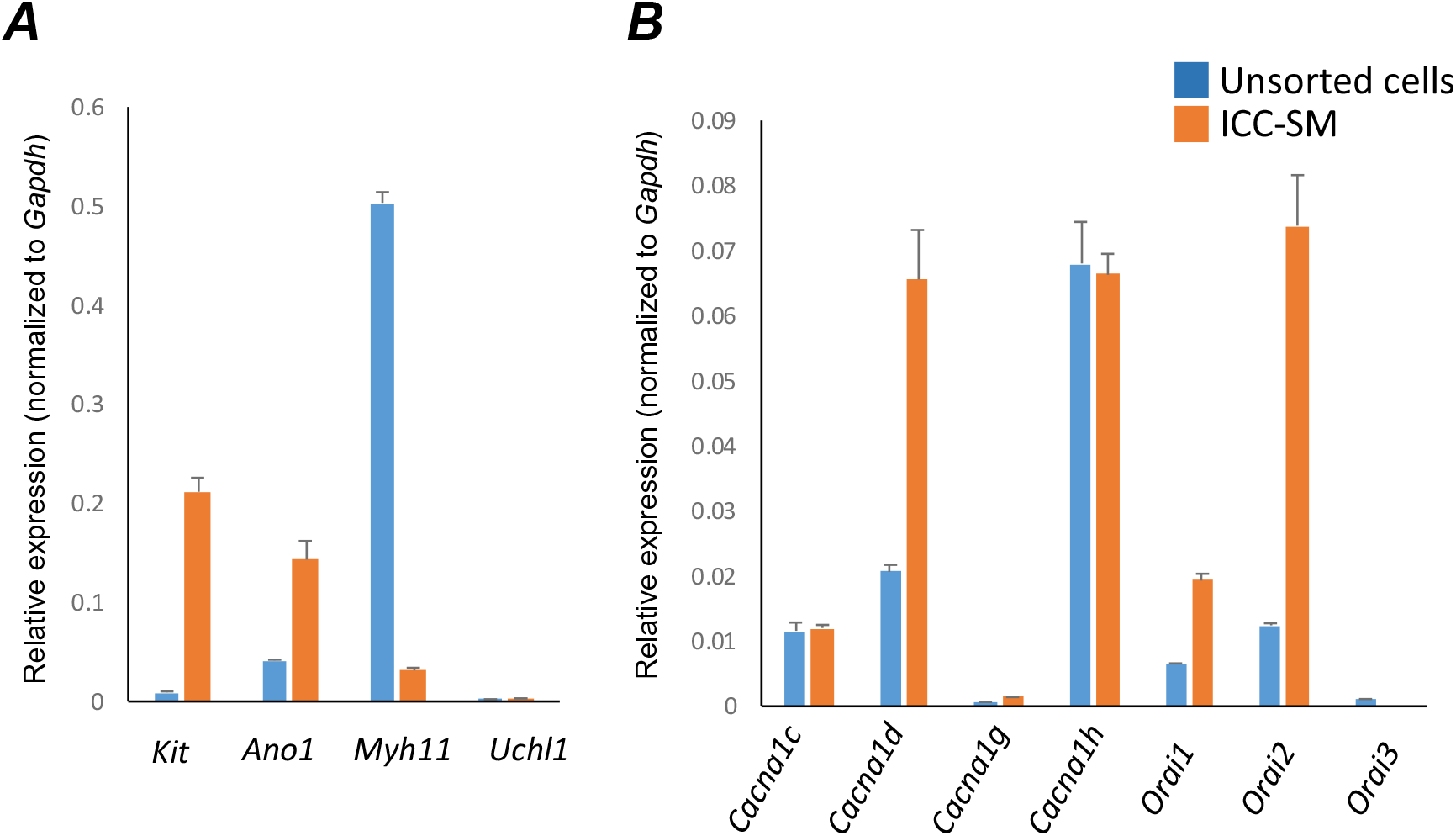
Molecular expression of genes related to Ca^2+^ entry channels. ***A*** Relative expression of cellular-specific biomarker genes in ICC-SM (sorted to purity by FACS) and compared with unsorted cells dispersed from submucosal tissues obtained from Kit^+*/copGFP*^ mice. Relative expression was determined by qPCR and normalized to *Gapdh* expression. Genes examined were *Kit* (tyrosine kinase receptor, found in ICC), Ano1 (Ca^2+^-activated Cl^-^ channel), *Uch11* (neural marker encoding PGP 9.5), *Myh11* (smooth muscle myosin). ***B*** Relative expression of major Ca^2+^ entry channels considered most likely to be expressed in colonic ICC from RNAseq of total ICC in murine colon [103]. L-Type Ca^2+^ channels (*Cacna1c* and *Cacna1d*) T-type Ca^2+^ channels (*Cacna1g* and *Cacna1h*) and Store-operated Ca^2+^ entry (SOCE) channels (*Orai1*, *Orai2* and *Orai3*) were evaluated. All data graphed as mean ± SEM (*n* = 4).

ICC-SM expressed L-type voltage-dependent Ca^2+^ channels encoded by *Cacna1c* and *Cacna1d* abundantly (Ca_V_ 1.2 and Ca_V_ 1.3 channels, respectively). *Cacna1c* showed a 0.012 ± 0.0005 and *Cacna1d* showed 0.066 ± 0.008 relative to *Gapdh* (Fig. 8 *B*; *n* = 4). ICC-SM also expressed *Cacna1h* (Ca_V_ 3.2) and to a lesser extent *Cacna1g* (Ca_V_ 3.1), both T-type voltage-dependent Ca^2+^ channels (Fig. 8 *B*; *n* = 4). *Cacna1h* expression was abundant in ICC-SM (0.07 ± 0.003 relative to *Gapdh*)*. Cacna1g* expression was less than *Cacna1h* 0.0014 ± 0.0001 relative to *Gapdh* (Fig. 8 *B*; *n* = 4).

Maintenance and refilling of cellular Ca^2+^ stores may also be important for mediation and shaping of Ca^2+^ signals in ICC-SM. Store-operated Ca^2+^ entry (SOCE) has been established as a mechanism for filling stores upon depletion [47, 48]. Contributions from SOCE via STIM and Orai interactions are essential for maintenance of Ca^2+^ stores in other types of ICC [49–52]. Colonic ICC-SM showed enrichment in *Orai1* and *Orai2* but *Orai3* was not resolved (Fig. 8 *B*; *n* = 4). *Orai1 expression* was 0.02 ± 0.001 relative to *Gapdh* and *Orai2* was 0.074 ± 0.008 relative to *Gapdh* (Fig. 8 *B*; *n* = 4).

### L-type Ca^2+^ channels are important for organization of Ca^2+^ transients in ICC-SM

L-type Ca^2+^ channels (Ca_V_ 1.3, Ca_V_ 1.2) were expressed in ICC-SM, and Ca^2+^ transients showed dependence on the Ca^2+^ gradient. Previous studies have reported that L-type channel antagonists inhibit slow waves in the colon [35, 53]. Therefore, we evaluated the contributions of Ca^2+^ entry via L-type channels to Ca^2+^ transients in ICC-SM. Nicardipine (1μM) abolished Ca^2+^ transients in ICC-SM (Fig. 9 *A&B*; *n* = 8). Firing site occurrence maps (Fig. 9 *C&D*; *n* = 8) describe the inhibitory effects of nicardipine on Ca^2+^ transients. Ca^2+^ PTCL area was reduced to 10.5 ± 4.7% (Fig. 9 *E-G*; *n* = 8) and PTCL count was reduced to 12.3 ± 4.8% (Fig. 9 *H*; *n* = 8). The number of firing sites also decreased to 8.4 ± 3% in the presence of nicardipine (Fig. 9 *I*; *n* = 8). Isradipine inhibits Ca_V_ 1.2 and Ca_V_ 1.3 equally [54, 55]. Isradipine (1μM) also reduced Ca^2+^ transients in ICC-SM (Supplemental Fig. 1 *A&B; n* = 7). ICC-SM Ca^2+^ firing was reduced in the presence of isradipine as shown in the firing sites occurrence maps (Supplemental Fig. 1 *C&D*). PTCL area was reduced to 18 ± 5 % (Supplemental Fig. 1 *E-G*; *n* = 7), and PTCL count was reduced to 19.5 ± 6 % (Supplemental Fig. 1 *H*; *n* = 7). The number of firing sites was inhibited by isradipine to 21.5 ± 6% (Supplemental Fig. 1 *I*; *n* = 7). These data suggest that ICC-SM Ca^2^ transients depend on Ca^2+^ influx via L-type Ca^2+^ channels and tend to suggest that Ca_V_ 1.2 is the dominant Ca^2+^ entry channel.

**Figure 9.**
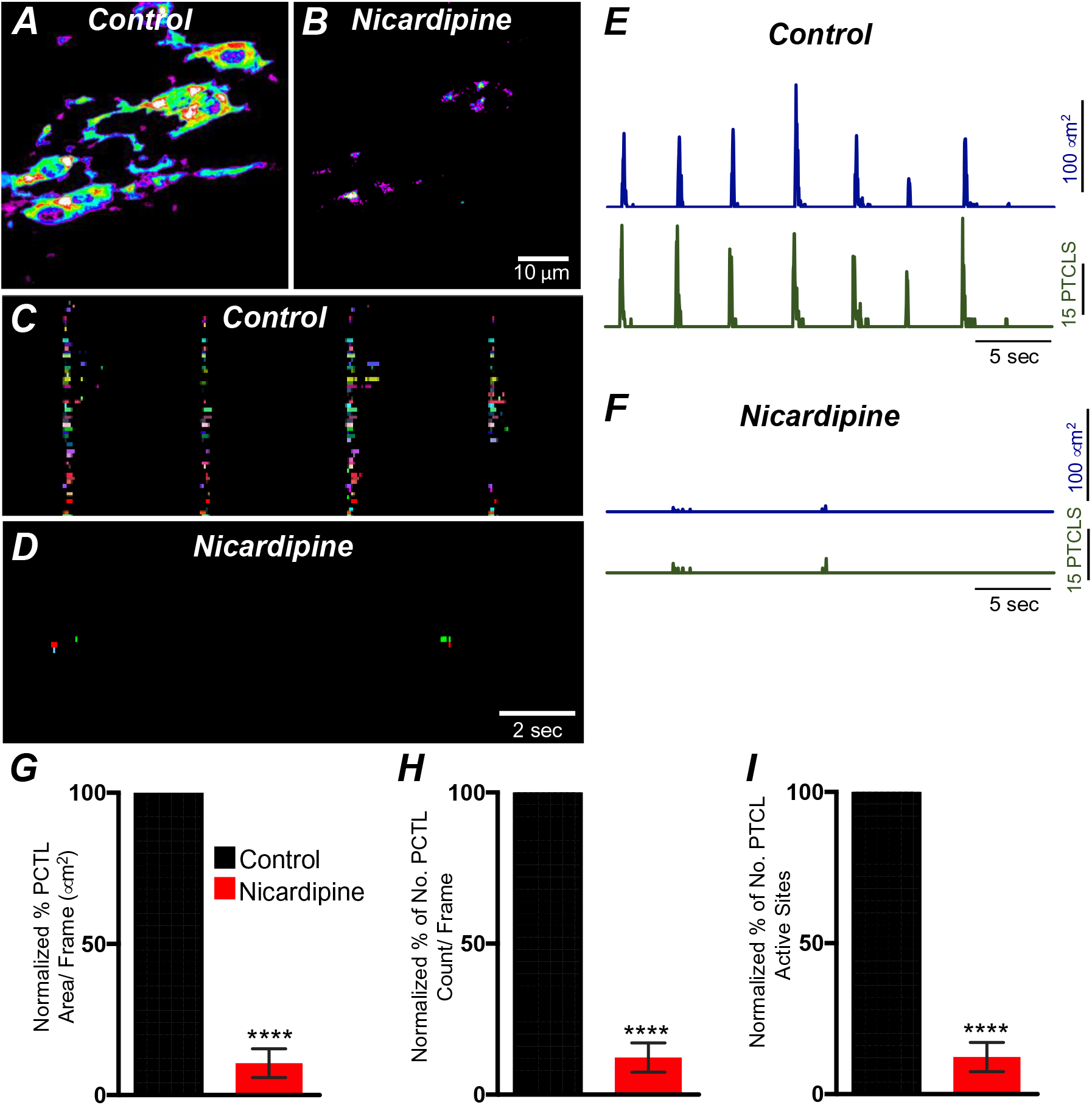
L-type Ca^2+^ channel antagonist, nicardipine effects on ICC-SM Ca^2+^ transients. ***A&B*** Representative heat-map images of an ICC-SM network from proximal colon of a Kit-iCre-GCaMP6f mouse showing active Ca^2+^ PTCLs under control conditions and in the presence of nicardipine (1μM). Ca^2+^ activity is color-coded with warm areas (white, red) representing bright areas of Ca^2+^ fluorescence and cold colors (purple, black) representing dim areas of Ca^2+^ fluorescence. Scale bar is 10 μm in both ***A&B***. ***C&D*** Firing sites showing Ca^2+^ activity in ICC-SM. Firing sites were color-coded and plotted as an occurrence map under control conditions and in the presence of nicardipine (1μM). Traces of firing sites showing PTCL area (***E***; blue) and PTCL count (***E***; green) under control conditions and in the presence of nicardipine; PTCL area (***F***; blue) and PTCL count (***F***; green) show the inhibitory effects of nicardipine on Ca^2+^ transients in ICC-SM. Summary graphs of Ca^2+^ PTCL activity in ICC-SM before and in the presence of nicardipine are shown in ***G*** (PTCL area/frame), ***H*** (PTCL count/frame) and the number of PTCL active sites ***I***. Data were normalized to controls and expressed as percentages (%). Significance determined using unpaired t-test, **** = *P*<0.0001, *n*= 8. All data graphed as mean ± SEM.

### T-type Ca^2+^ channels also contribute to Ca^2+^ transients in ICC-SM

T-type Ca^2+^ channels are an important source for Ca^2+^ entry in pacemaker ICC of the small intestine [56–58]. T-type channel antagonists (Ni^2+^ and mibefradil) reduced the rate-of-rise of the upstroke component of the slow waves and higher concentrations attenuated slow wave activity [56, 59, 60]. As reported above, ICC-SM express T-type channels *Cacna1h* and *Cacna1g.* Therefore, the role of T-type Ca^2+^ channels in modulating Ca^2+^signaling in ICC-SM was evaluated with the specific T-type channel antagonists NNC 55-0396 (10 μM), TTA-A2 (10 μM) and Z-944 (1 μM). NNC 55-0396 reduced Ca^2+^ transient firing (Fig. 10 *A&B*), Ca^2+^ transients firing sites occurrence maps (Fig. 10 *C&D*) and firing sites. PTCL area and PTCL count traces show a reduction in Ca^2+^ transient firing (Fig. 10 *E&F*). PTCL area was reduced to 31.7 ± 3.3 % (Fig. 10 *E-G*; *n* = 9) and PTCL count was reduced to 35.6 ± 4.5 % (Fig. 10 *H*; *n* = 9). The number of firing sites was also reduced by NNC 55-0396 to 37.6 ± 4.3 % (Fig. 10 *I*; *n* = 9). TTA-A2 showed similar inhibitory effects on ICC-SM Ca^2+^ transients (Supplemental Fig. 2). PTCL area was reduced to 42 ± 5.8 % (Supplemental Fig. 2 *E*; *n* = 7) and PTCL count was reduced to 44.4 ± 6.2 % (Supplemental Fig. 2 *F*; *n* = 7) The number of firing sites was also reduced by TTA-A2 to 43.4 ± 6 % (Supplemental Fig. 2 *G*; *n* = 7). Z-944 significantly reduced ICC-SM Ca^2+^ transients but was somewhat less effective than NNC 55-0396 or TTA-A2. PTCL area was reduced to 56 ± 10 % (Supplemental Fig. 2 *E*; *n* = 5), and PTCL count was reduced to 53 ± 12 % (Supplemental Fig. 2 *F*; *n* = 5) The number of firing sites was also reduced by Z-944 to 53.4 ± 11 % (Supplemental Fig. 2 *G*; *n* = 5). The data suggest that T-type Ca^2+^ channels contribute to the initiation and organization of ICC-SM Ca^2+^ firing.

**Figure 10.**
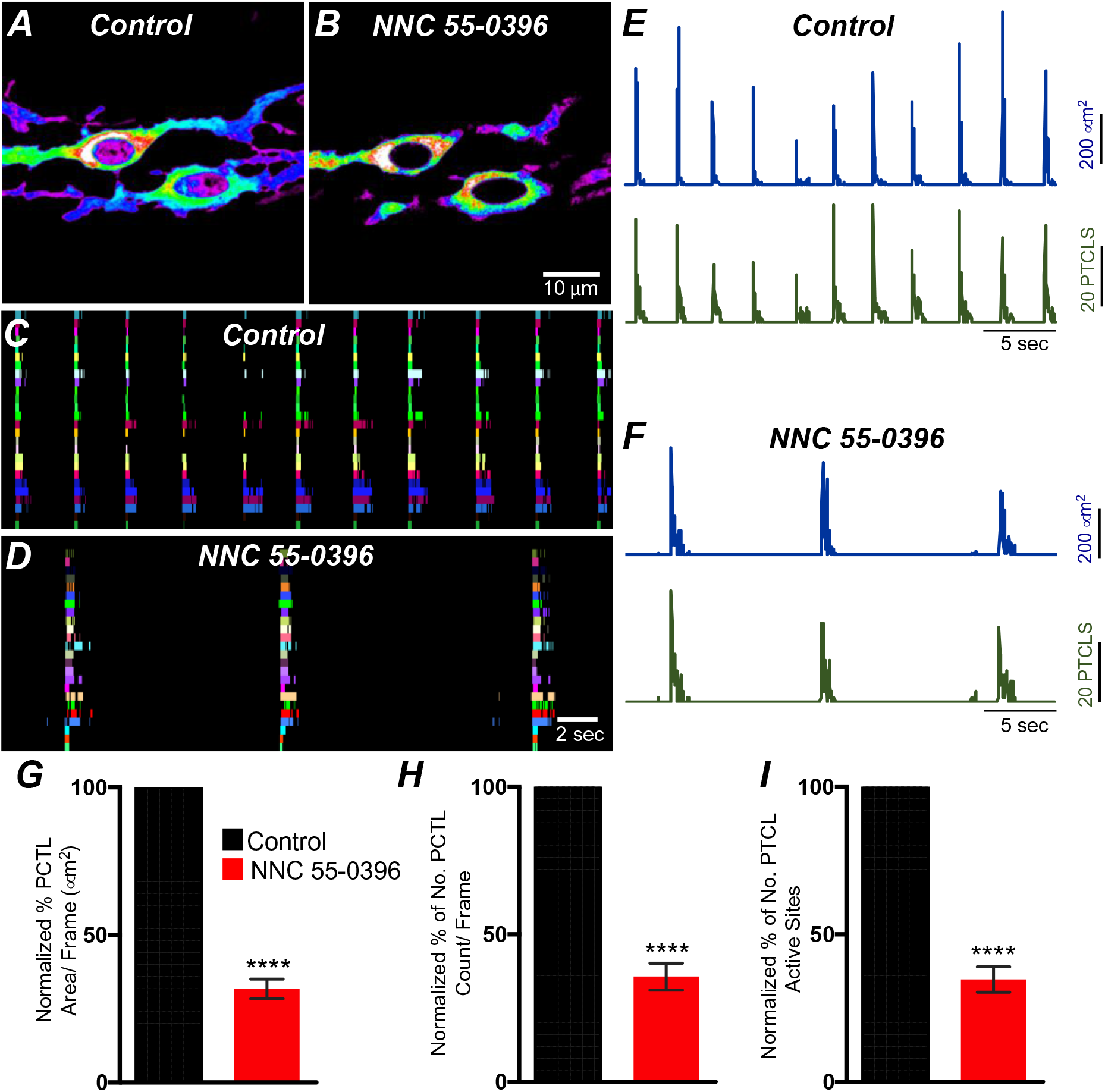
T-type Ca^2+^ channel antagonist, NNC 55-0396 effects on ICC-SM Ca^2+^ transients. ***A&B*** Representative heat-map images of Ca^2+^ transient particles in ICC-SM under control conditions ***A*** and in the presence of NNC 55-0396 (10 μM) ***B***. Active firing sites were color-coded and plotted as an occurrence maps in the ICC-SM network under control ***C*** and NNC 55-0396 ***D*** conditions. Plots of Ca^2+^ transient particle activity of ICC-SM in control conditions and in the presence of NNC 55-0396 showing PTCL area (blue) and PTCL count (green) under control conditions ***E*** and in the presence of NNC 55-0396 ***F***. Summary graphs of average percentage changes in PTCL area ***G***, PTCL count ***H*** and the number of Ca^2+^ firing sites after addition of NNC 55-0396 ***I***. Data were normalized to controls and expressed as percentages (%). Significance determined using unpaired t-test, **** = *P*<0.0001, *n*= 8. All data graphed as mean ± SEM.

### Effects of membrane hyperpolarization on Ca^2+^ transients in ICC-SM

Experiments described above showed that Ca^2+^ transients in ICC-SM depend on voltage-dependent Ca^2+^ influx mechanisms (Fig. 7-10). The role and contributions of the voltage-dependent channels expressed in ICC-SM (Ca_V_ 1.3 and Ca_V_ 1.2 and Ca_V_ 3.2) was further examined under conditions of membrane hyperpolarization induced by activation of K_ATP_ channels. K_ATP_ is functional in colonic SMCs but not in ICC [61]. Therefore, for these experiments Ca^2+^ imaging was performed on preparations in which the submucosal layer remained attached to the muscularis. Pinacidil (10 μM), a selective K_ATP_ channel agonist, hyperpolarizes murine colonic muscles by ∼ 10 mV [56, 62, 63]. Pinacidil showed a tendency toward increased Ca^2+^ transient firing, but this effect did not reach statistical significance (Fig. 11 *A-I*). Ca^2+^ PTCL area was increased to 130.3 ± 24.3% (Fig. 11 *G*; *P value =0.24; n* = 6) and PTCL count was increased to 121.8 ± 17.8% (Fig. 11 *H*; *P value =0.25; n* = 6). The number of firing sites was not affected by pinacidil (Fig. 11 *I*; *P value =0.22; n* = 6). The effects of pinacidil on global Ca^2+^ transients were evaluated. Pinacidil significantly reduced the duration of global Ca^2+^ transients from 1.78 ± 0.2 s to 0.89 ± 0.1 s (Fig. 11 *J&K*; *n* = 6). Reduction in the duration of Ca^2+^ transients was associated with a tendency for an increase in frequency, but this parameter did not reach significance. Ca^2+^ oscillation frequency under control conditions was 15.1 ± 1.1 cpm and in the presence of pinacidil was 16.5 ± 1.2 cpm (Fig. 11 *J&L*; *P value =0.45; n* = 6).

**Figure 11.**
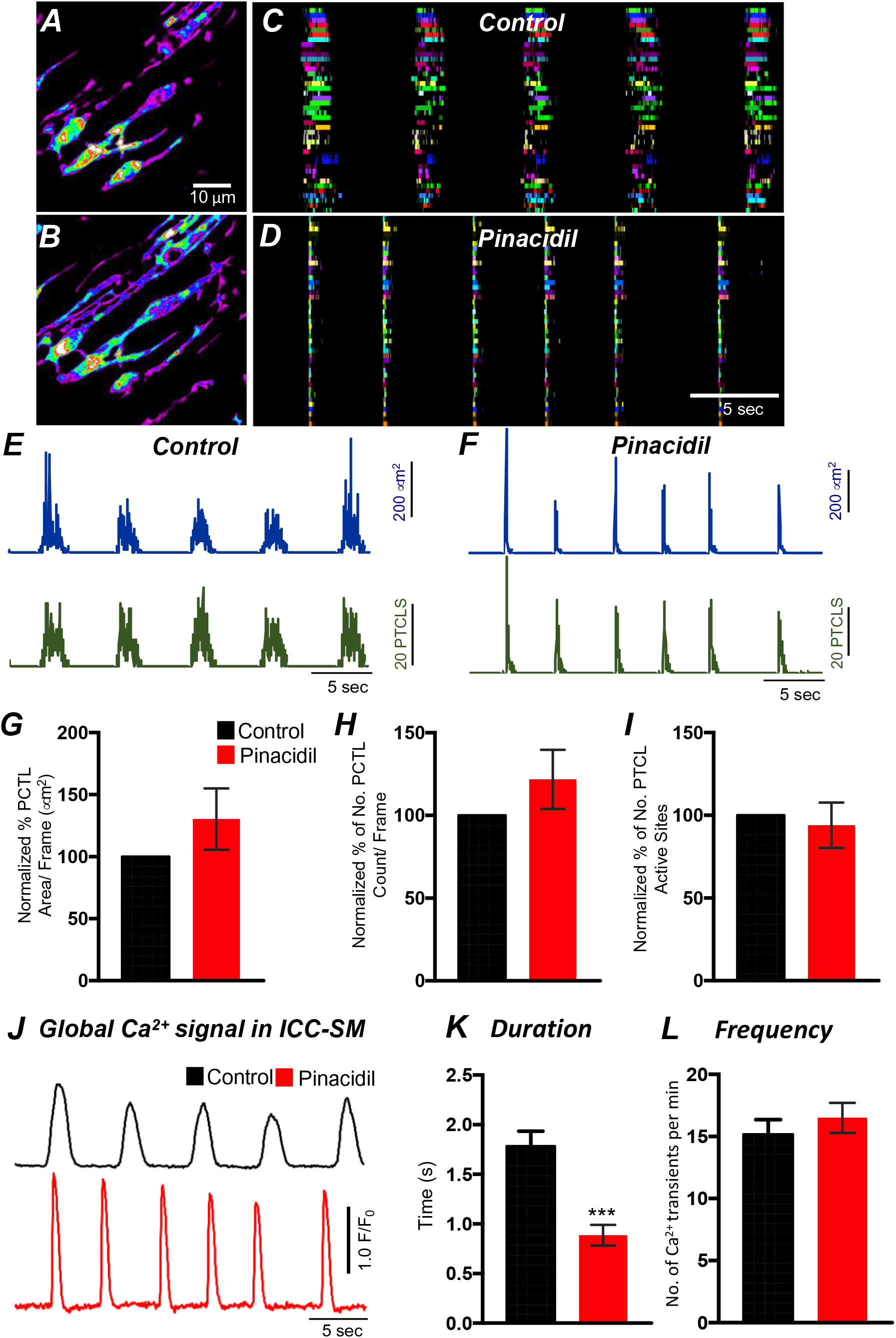
The effects of pinacidil of Ca^2+^ transients in ICC-SM. ***A*** Representative heat-map image of an ICC-SM network showing total Ca^2+^ PTCLs under control conditions and in the presence of pinacidil (10 μM) ***B***. Ca^2+^ activity was color-coded with warm areas (white, red) representing bright areas of Ca^2+^ fluorescence and cold colors (purple, black) representing dim areas of Ca^2+^ fluorescence. Scale bar is 10 mm in both ***A&B***. ***C&D*** Ca^2+^ firing sites in ICC-SM were color-coded and plotted as occurrence maps under control conditions in ***C*** and in the presence of pinacidil (10 μM) ***D***. Traces of firing sites PTCL area (***E***; blue) and PTCL count (***E***; red) under control conditions and in the presence of pinacidil PTCL area (***F***; blue) and PTCL count (***F***; red). Summary graphs of Ca^2+^ PTCL activity in ICC-SM in the presence of pinacidil are shown in ***G*** (PTCL area), ***H*** (PTCL count) and the number of PTCL active sites ***I***. Global Ca^2+^ traces are plotted from all cells in the FOV in ***B*** under control conditions (black trace) and in the presence of pinacidil (red trace). ***K&L*** Summary graphs of the duration (s) ***K*** and frequency (cpm) ***L*** of global calcium transients in ICC-SM. Data were normalized to controls and expressed as percentages (%) in G,H and I. Significance determined using unpaired t-test, *** = *P*<0.001, *n*= 6. All data graphed as mean ± SEM.

The effects of nicardipine were tested in the presence of pinacidil. Under these conditions nicardipine significantly reduced Ca^2+^ transients in ICC-SM (Fig. 12 *G&H*; *n* = 5). PTCL area was reduced to 38.5 ± 7.6 % (Fig. 12 *G*; *n* = 5) and PTCL count was reduced to 42.3 ± 9.1 % (Fig. 12 *H*; *n* = 5). In some regards these results were surprising as membrane potential hyperpolarization might reduce contributions from L-type Ca^2+^ channels (Ca_V_ 1.2). One explanation is that Ca_V_ 1.3, which are abundant in ICC-SM and activate at more negative potentials than Ca_V_ 1.2 [64] may contribute to Ca^2+^ entry at more hyperpolarized potentials. Further addition of NNC 55-0396 (10 μM) inhibited Ca^2+^ transients in ICC-SM to a greater extent. PTCL area was reduced to 9.0 ± 2 % (Fig. 12 *G*; *n* = 5), and PTCL count was reduced to 11.2 ± 1.9 % (Fig. 12 *H*; *n* = 5). The utilization of two Ca^2+^ conductances with different ranges of voltage-dependent activation for the initiation of CTCs provides a safety factor that insures persistence of pacemaker activity over a broad range of membrane potentials.

**Figure 12.**
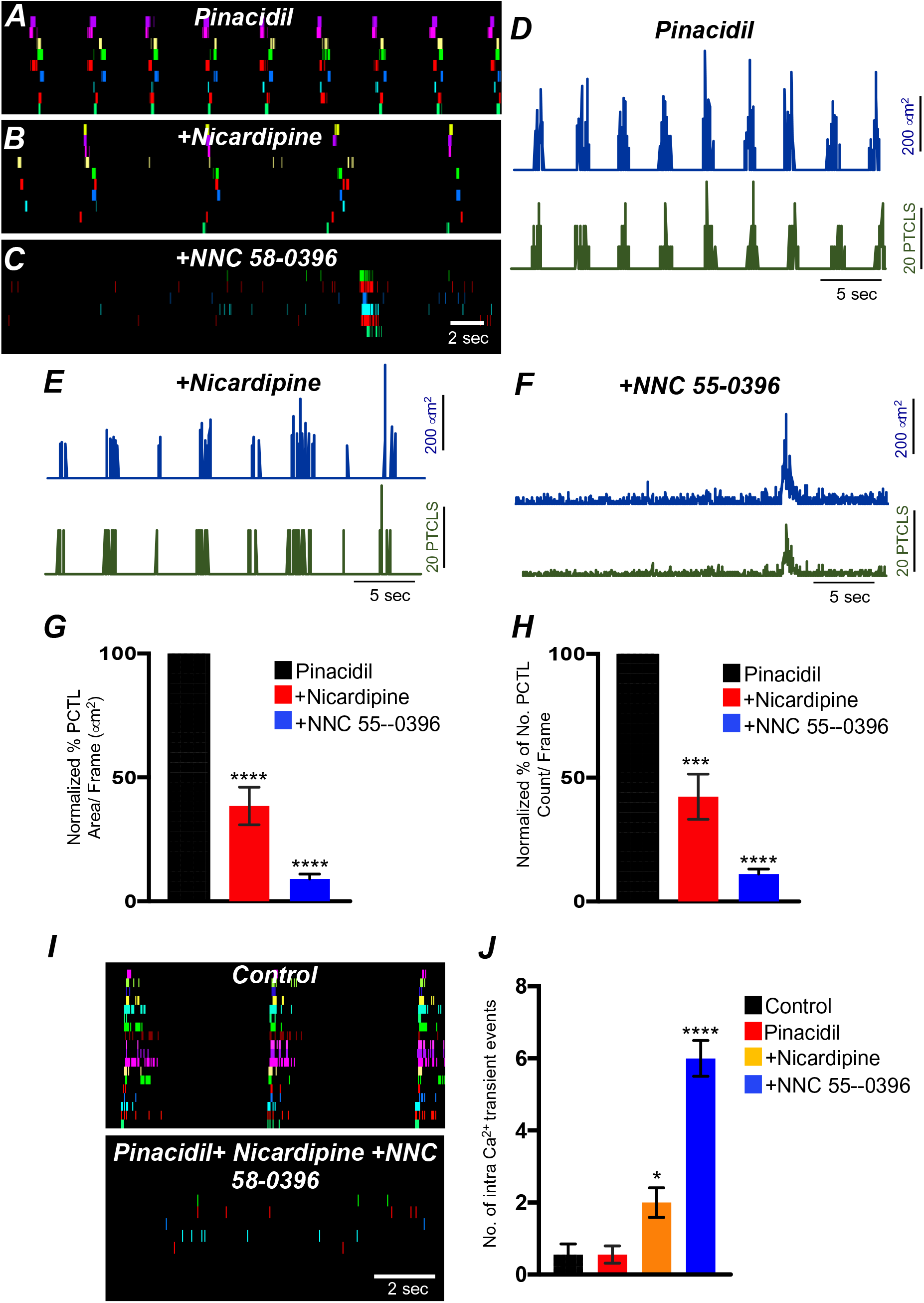
The effects of membrane hyperpolarization and voltage-dependent Ca^2+^ entry block on ICC-SM Ca^2+^ transients. ***A*** Ca^2+^ Firing sites in ICC-SM are color-coded and plotted in occurrence map in the presence of pinacidil (10 μM). ***B*** Shows effects of nicardipine (1 μM) in the continued presence of pinacidil. ***C*** shows effects of combining nicardipine and NNC 55-0396 (10 μM) in the continued presence of pinacidil. Traces of firing sites PTCL area (blue) and PTCL count (green) under each condition are shown in ***D*** pinacidil ***E*** pinacidil & nicardipine and ***F*** combination of pinacidil, nicardipine and NNC 55-0396. Summary graphs of Ca^2+^ PTCL activity in ICC-SM in the presence of voltage dependent Ca^2+^ channel antagonists (nicardipine and NNC 55-0396) are shown in ***G*** (PTCL area) and ***H*** (PTCL count). ***I*** The number of Ca^2+^ firing events were tabulated during 2s intervals before the initiation of Ca^2+^ transient clusters in ICC-SM (period of tabulation indicated by the grey box) and summarized in ***J*** under control conditions, in pinacidil, in pinacidil & nicardipine and in a combination of pinacidil, nicardipine and NNC 55-0396. Data were normalized to controls and expressed as percentages (%) in **G** and **H**. Significance determined using one-way-ANOVA, * = *P*<0.1, ** = *P*<0.01, *** = *P*<0.001 **** = *P*<0.0001, *n*= 5. All data graphed as mean ± SEM.

Inhibiting voltage dependent Ca^2+^ channels in the presence of pinacidil unmasked underlying Ca^2+^ transients that occurred more randomly than the clustered transients occurring normally (Fig. 12*F&I*). We tabulated the number of Ca^2+^ events in the intervals between CTCs (calculated from a period of 2s before the onset a CTC). Underlying Ca^2+^ events were more frequent in the presence of pinacidil and nicardipine and increased again upon addition of NNC 55-0396 (Fig. 12 *J*; *n* = 5).

### Contributions of intracellular Ca^2+^ stores and release channels in ICC-SM Ca^2^ activity

Previous studies have demonstrated that Ca^2+^ signaling in ICC-MY in the small intestine depends not only on Ca^2+^ influx but also on Ca^2+^ release from intracellular stores [44]. Ca^2+^ release from stores is also critical for generation of pacemaker currents and slow waves [65–67]. The role of Ca^2+^ release mechanisms in Ca^2+^ signaling in ICC-SM was also evaluated. Thapsigargin (1 μM; A SERCA pump antagonist) reduced, but did not block, Ca^2+^ transient firing in ICC-SM (Fig. 13 *A-D*). PTCL area was reduced to 29 ± 12% (Fig. 13 *E*; *n* = 6) and PTCL count was reduced to 22 ± 10.1% (Fig. 13 *F*; *n* = 6). The number of firing sites was also reduced by thapsigargin to 21 ± 11% (Fig. 13 *G*; *n* = 6). Cyclopiazonic acid (CPA, 10 μM), another SERCA antagonist, also reduced Ca^2+^ transient firing (Fig. 13 *H-K*). PTCL area was reduced to 36.1 ± 8.3% (Fig. 13 *L*; *n* = 5) and PTCL count was reduced to 29.1 ± 6.3% (Fig. 13 *M*; *n* = 5). The number of firing sites was reduced by CPA to 56 ± 9.5 % (Fig. 13 *N*; *n* = 5).

**Figure 13.**
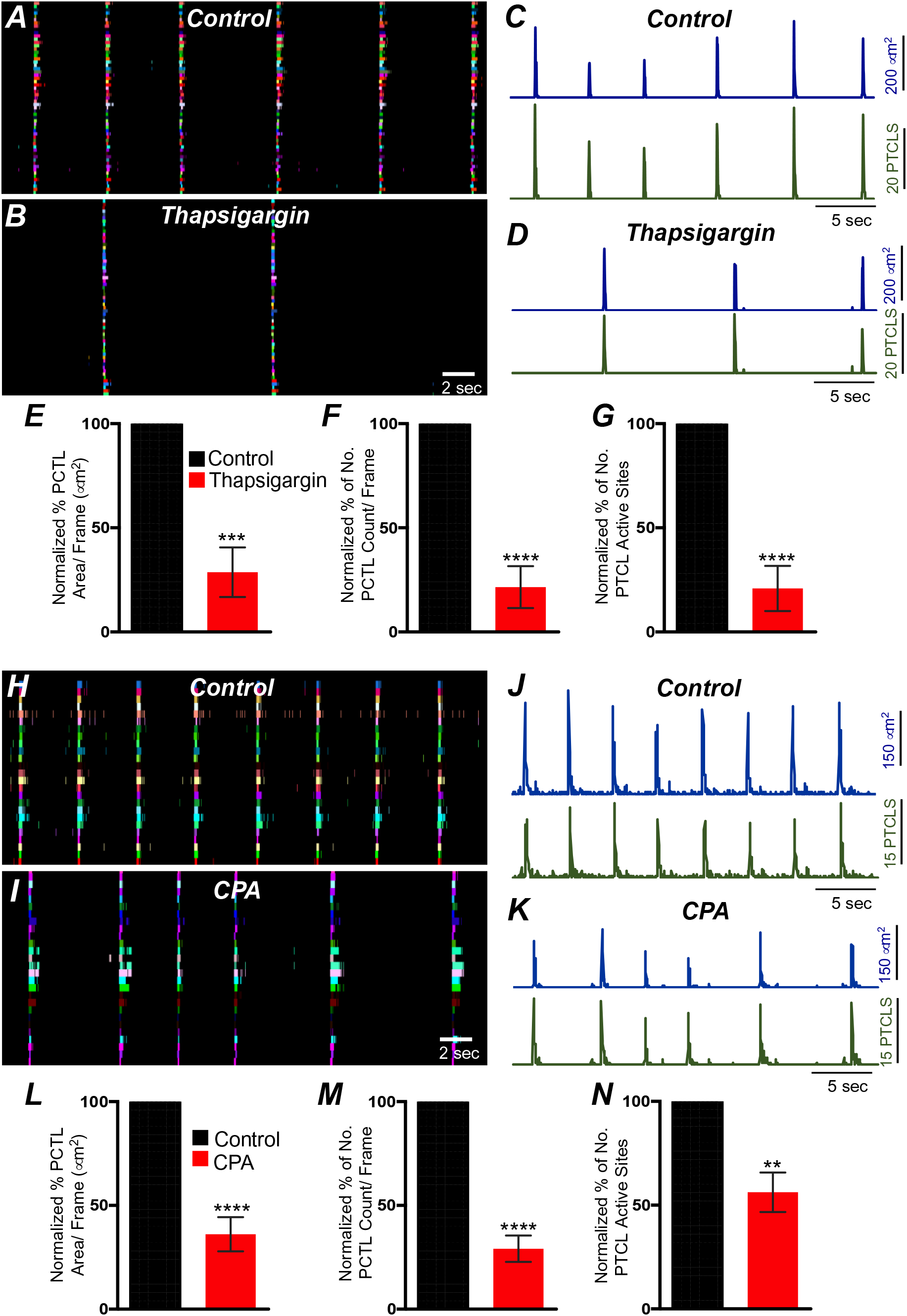
Contributions of Ca^2+^ stores in overall ICC-SM Ca^2+^ transients. ***A*** Ca^2+^ activity of firing sites in ICC-SM are color-coded and plotted in occurrence maps under control conditions and in the presence of thapsigargin (1 μM) ***B***. Traces of firing sites PTCL area (***C***; blue) and PTCL count (***C***; green) under control conditions and in the presence of thapsigargin PTCL area (***D***; blue) and PTCL count (***D***; green). Scale bars in ***C*** applies to traces in D. Summary graphs of Ca^2+^ PTCL activity in ICC-SM in the presence of thapsigargin are shown in ***E*** (PTCL area), ***F*** (PTCL count) and the number of PTCL active sites ***G*** (*n*= 6). CPA (SERCA pump inhibitor) reduced transients compared to control as shown in occurrence maps of firing sites ***H&I*** and Ca^2+^ activity traces ***J&K.*** Scale bars in J applies to traces in K. Summary graphs of Ca^2+^ PTCL activity in ICC-SM in the presence of CPA are shown in ***L*** (PTCL area), ***M*** (PTCL count) and the number of PTCL active sites ***N*** (*n*= 5). Data were normalized to controls and expressed as percentages (%). Significance determined using unpaired t-test, ** = *P*<0.01, *** = *P*<0.001 **** = *P*<0.0001. All data graphed as mean ± SEM.

ER release channels RyRs and IP_3_Rs amplify and sustain Ca^2+^ signaling via direct localized release of Ca^2+^ or Ca^2+^-induced Ca^2+^ release (CICR) [68–75]. Contributions from RyRs and IP_3_Rs to Ca^2+^ transients in ICC-SM were therefore investigated. Ryanodine (100 μM) significantly reduced Ca^2+^ event firing in ICC-SM (Fig. 14 *A-F*). PTCL area was reduced to 76.1 ± 1.6 % (Fig. 14 *G*; *n* = 4) and PTCL count was reduced to 80.9 ± 2.7 % (Fig. 14 *H*; *n* = 4). The number of firing sites was also reduced by ryanodine to 75.7 ± 2.1 % (Fig. 14 *I*; *n* = 4). We also noted that the greatest effects of ryanodine on Ca^2+^ transients occurred after the first ∼300 ms of CTCs (Fig. 14 *J&K*; *n* = 4). Thus, ryanodine shortens the durations of the CTCs. Ca^2+^ release mechanisms via RyRs contribute to the overall patterns of Ca^2+^ waves in ICC-SM, as shown by the distribution plots of average percentages of firing sites during a Ca^2+^ wave (Fig. 14 *K*; *n* = 4).

**Figure 14.**
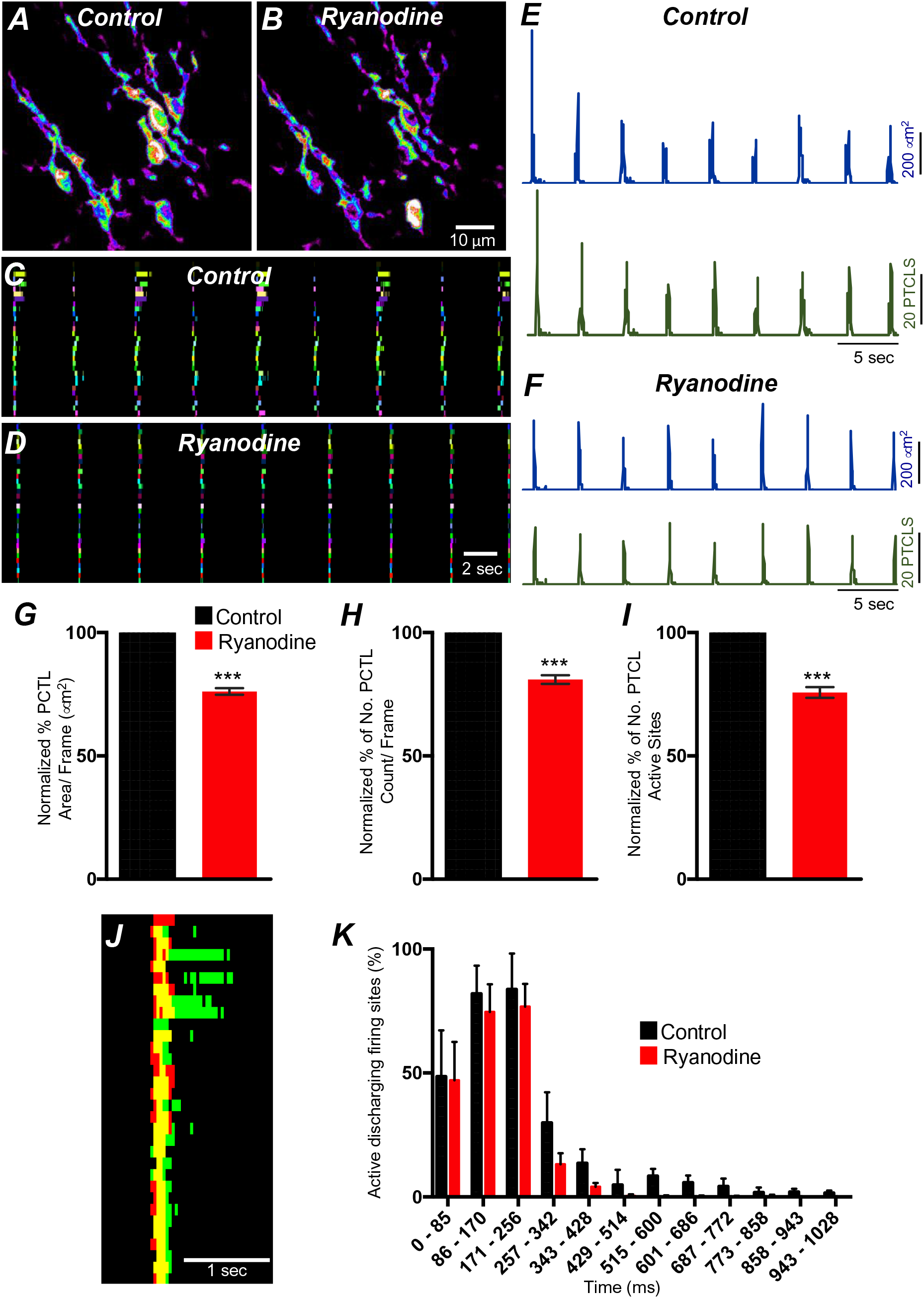
Ryanodine receptors (RyRs) contributing to Ca^2+^ release in ICC-SM. ***A*** Representative heat-map image of an ICC-SM network from proximal colon showing total active Ca^2+^ PTCLs under control conditions and in the presence of ryanodine (100 μM) ***B***. ***C&D*** Ca^2+^ firing sites are color-coded and plotted in occurrence maps showing the effect of the ryanodine (100 μM), on Ca^2+^ transient clusters (CTCs) in ICC-SM. Traces of firing sites PTCL area (***E***; blue) and PTCL count (***E***; green) under control conditions and in the presence of ryanodine, PTCL area (***F***; blue) and PTCL count (***F***; green). Summary graphs of Ca^2+^ PTCL activity in ICC-SM in the presence of ryanodine are shown in ***G*** (PTCL area), ***H*** (PTCL count) and the number of PTCL active sites ***I*** (*n*= 4). ***J*** overlaid occurrence maps of showing Ca^2+^ firing during control conditions (all firing sites are in green) and in the presence of ryanodine (all firing sites are in red). Note how ryanodine shortened the duration of the Ca^2+^ transient cluster (CTC). ***K*** Distribution plot of average percentages of firing sites during a Ca^2+^ wave, calculated for 1 s duration and plotted in 85 ms bins showing that ryanodine mainly blocked Ca^2+^ transients occurring after the first 257 ms intervals (*n*= 4). Data were normalized to controls and expressed as percentages (%). Significance determined using unpaired t-test, *** = *P*<0.001. All data graphed as mean ± SEM.

Xestospongin C (10 μM; An IP_3_R antagonist) also reduced Ca^2+^ events in ICC-SM (Fig. 15 *A-I*; *n* = 4). PTCL area was significantly reduced to 78 ± 5.8% (Fig. 15 *G*; *n* = 4) and although both PTCL count and number were reduced to 75.3 ± 10.5% and 74.8 ± 9.3%, respectively, these effects did not reach statistical significance (PTCL count: P value = 0.08 and PTCL number: P value = 0.06; Fig. 15 *H&I*; *n* = 4). Xestospongin C displayed inhibitory effects similar to ryanodine; most of the inhibition of Ca^2+^ events occurred after the first ∼400 ms of CTCs (Fig. 15 *J&K*; *n* = 4). Thus, Ca^2+^ release via IP_3_Rs also contributes to the overall pattern of Ca^2+^ waves in ICC-SM, as shown by the distribution plots of average percentages of firing sites during a Ca^2+^ wave (Fig. 15 *K*; *n* = 4).

**Figure 15.**
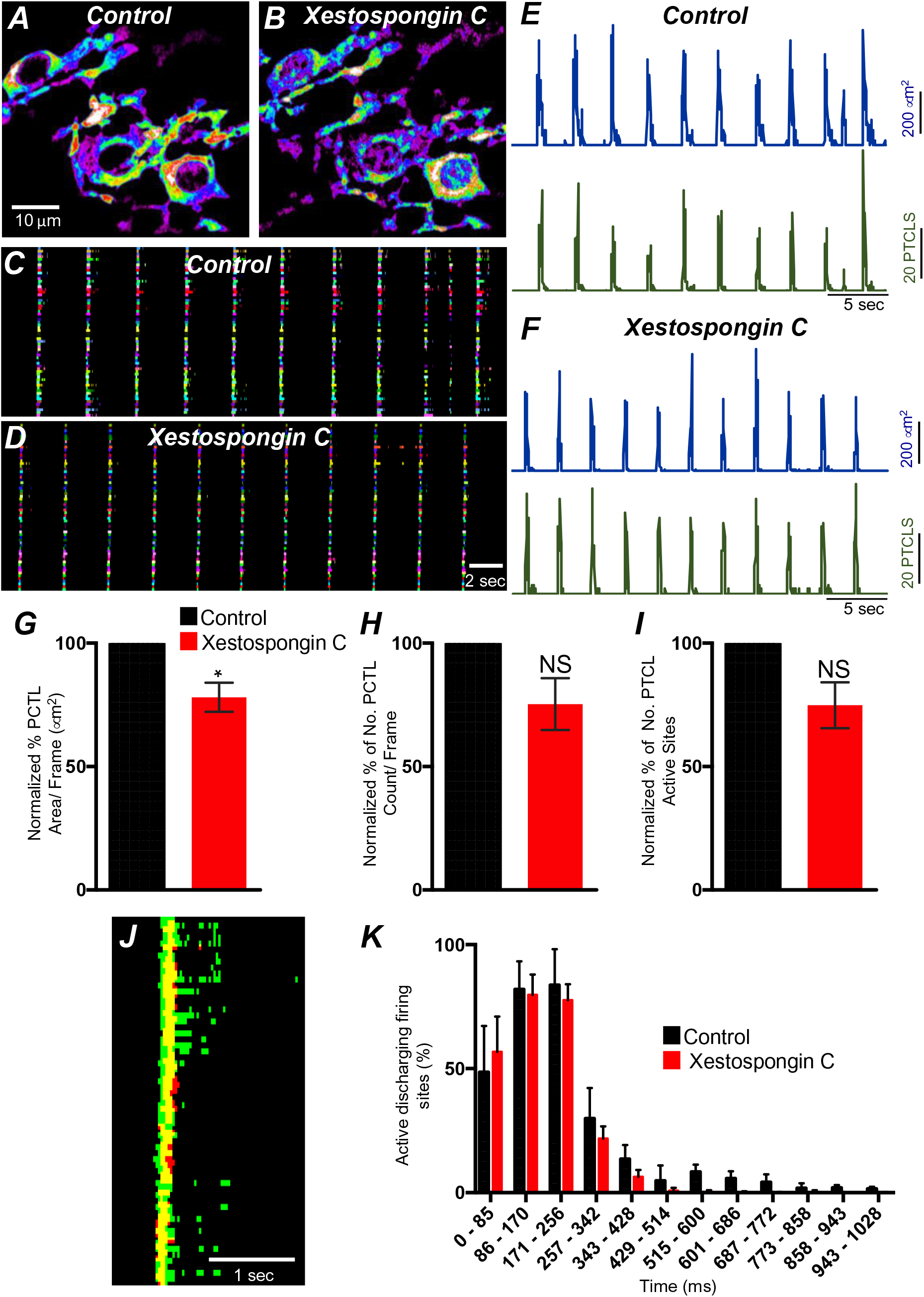
IP_3_ receptors (IP_3_Rs) contribution to Ca^2+^ transients in ICC-SM. ***A&B*** Representative images of heat-maps of the summated Ca^2+^ transient particles of ICC-MY under control conditions ***A*** and in xestospongin C (10 μM; ***B***). ***C&D*** Ca^2+^ firing sites are color-coded and plotted in occurrence maps showing the effect of the xestospongin C (1 μM) on Ca^2+^ transient clusters (CTCs). Traces of firing sites PTCL area (***E***; blue) and PTCL count (***E***; green) under control conditions and in the presence of xestospongin C, PTCL area (***F***; blue) and PTCL count (***F***; green). Summary graphs of Ca^2+^ PTCL activity in ICC-SM in the presence of xestospongin C are shown in ***G*** (PTCL area), ***H*** (PTCL count) and the number of PTCL active sites ***I*** (*n*= 4). ***J*** overlaid images of Ca^2+^ event firing in control (all firing sites are in green) and in the presence of xestospongin C (all firing sites are in red). Note how ryanodine shortened the duration of the CTC. ***K*** Distribution plot of average percentages of firing sites during a Ca^2+^ wave, calculated for 1 s duration and plotted in 85 ms bins showing that xestospongin C mainly blocked Ca^2+^ transients occurring after the 257-342 ms intervals (*n*= 4). Data were normalized to controls and expressed as percentages (%). Significance determined using unpaired t-test, * = *P*<0.1 and Not significant (NS) = *P*>0.05. All data graphed as mean ± SEM.

2-APB (100 μM) and tetracaine were also tested (100 μM; Supplemental Fig. 3 *A&B*; *n* = 5) as secondary tests of the contributions of IP_3_Rs and RyRs in CTCs. 2-APB reduced Ca^2+^ PTCL area to 61.1 ± 14.5% (Supplemental Fig. 3 *Ai*; *n* = 5) and reduced PTCL count to 58 ± 13.2% (Supplemental Fig. 3 *Aii*; *n* = 5). The number of firing sites was also reduced by 2-APB to 63.7 ± 11.3% (Supplemental Fig. 3 *Aiii*; *n* = 5). Tetracaine reduced Ca^2+^ PTCL area to 85 ± 5.1% (Supplemental Fig. 3 *Bi*; *n* =5) and reduced PTCL count to 69.5 ± 7.8 % (Supplemental Fig. 3 *Bii*; *n* = 5). The number of firing sites was also reduced by tetracaine to 71.3 ± 5.9% (Supplemental Fig. 3 *Biii*; *n* = 5).

The effects of 2-APB could be non-specific and may include effects on store-operated Ca^2+^ entry channels (SOCE; e.g. by blocking Orai channels). Previous studies have shown SOCE to be important for maintenance of Ca^2+^ stores and sustaining Ca^2+^ release from the ER [49, 76–80]. ICC-SM express Orai channels (*Orai1* and Orai*2*; Fig. 8 *B*), so the role of SOCE in maintenance of Ca^2+^ transients was examined using an Orai antagonist. GSK 7975A (10 μM; An Orai antagonist) reduced the firing frequency of CTCs (Fig. 16 *A&B*). Firing site occurrence (Fig. 16 *C&D*) and PTCL counts and areas were reduced (Fig. 16 *E&F*). Ca^2+^PTCL area was reduced to 42.4 ± 9.4 % (Fig. 16 *E-G*; *n* = 7) and PTCL count was reduced to 48 ± 7 % (Fig. 16 *H*; *n* = 7). The number of firing sites was also inhibited by GSK 7975A to 47.5 ± 4.1% (Fig. 16 *I*; *n* = 7).

**Figure 16.**
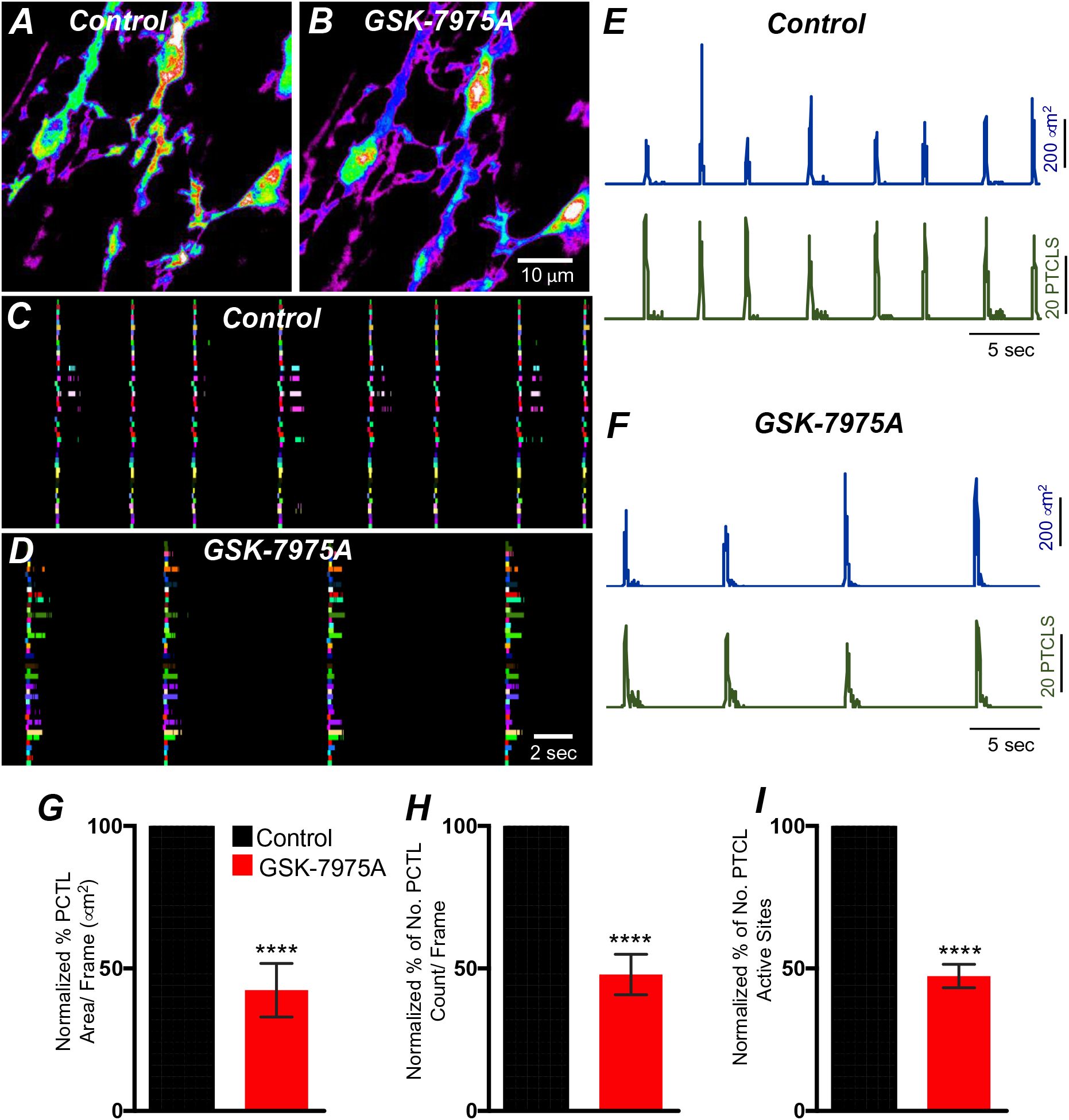
SOCE role in maintaining ICC-SM Ca^2+^ transients. ***A*** Representative Heat-map image of an ICC-SM network showing total active Ca^2+^ PTCLs under control conditions and in the presence of GSK-7975A (10 μM, for 20 min) ***B***. ***C&D*** Ca^2+^ firing sites are color-coded and plotted in occurrence maps showing the effect of the SOCE channel antagonist, GSK-7975A (100 μM), on ICC-SM Ca^2+^ transients. Traces of PTCL area (***E***; blue) and PTCL count (***E***; green) under control conditions and in the presence of GSK-7975A, PTCL area (***F***; blue) and PTCL count (***F***; green). Summary graphs of Ca^2+^ PTCL activity in ICC-SM in the presence of GSK-7975A are shown in ***G*** (PTCL area), ***H*** (PTCL count) and the number of PTCL active sites ***I*** (*n*= 7). Significance determined using unpaired t-test, **** = *P*<0.0001. All data graphed as mean ± SEM.

## Discussion

This study characterized Ca^2+^ transients responsible for the pacemaker function of ICC-SM that contributes to contractile patterns in colonic motility. The sequence of activation from ICC-SM to colonic SMCs was quantified using two optogenetic sensors, expressed specifically in ICC or SMCs. Correlation analysis demonstrated a 1:1 relationship between CTCs in ICC-SM and Ca^2+^ signaling and contractile responses in SMCs. The clustered Ca^2+^ transients in ICC-SM consisted of ∼2 sec bursts of activity from multiple sites within cells. Organization of the Ca^2+^ transients into clusters was due to voltage-dependent Ca^2+^ entry that appeared to be facilitated by both high-voltage activated Ca^2+^ channels (L-type encoded by *Cacna1c* and *Cacna1d* in ICC-SM) and low-voltage activated Ca^2+^ channels (T-type encoded in ICC-SM by *Cacna1h* and possibly *Cacna1g*). Part of the Ca^2+^ transients that made up CTCs was due to Ca^2+^ entry, as the earliest Ca^2+^ transients resolved within a cluster were not blocked by thapsigargin, CPA or antagonists of ryanodine and IP_3_ receptors. The earliest events were blocked by nicardipine, suggesting that L-type Ca^2+^ channels are the dominant Ca^2+^ entry pathway at basal resting potentials. Ca^2+^ transients in ICC-SM were not as sensitive to block by STIM and Orai, as in other ICC [49, 81], However, the Orai antagonist reduced the frequency of CTCs and may have blocked these events completely with longer treatment periods. Our data show that Ca^2+^ entry is fundamental in ICC-SM Ca^2+^ transients. Localized Ca^2+^ influx via clusters of L-type Ca^2+^ channels may result in localized elevations in [Ca^2+^]_i_ and activation of ANO1 channels directly. Localized Ca^2+^ entry through L-type Ca^2+^ channels has been termed Ca^2+^sparklets, and these events could be involved in Ca^2+^ influx [82–84] and initiation of pacemaker activity in ICC-SM. However, the decrease in the frequency of CTCs by manipulations to reduce store Ca^2+^ suggest an important role for Ca^2+^ release, perhaps resulting from coupling between Ca^2+^ entry and CICR [85] in regulating pacemaker activity.

In this study we developed a new preparation in which ICC-SM adherent to the submucosa was used to allow very high-resolution imaging without complications from muscular contractions. Preparations of this type may be valuable for future studies of cellular mechanisms responsible for pacemaker activity and factors that regulate or degrade pacemaker activity in pathophysiological conditions.

The pacemaker function of ICC-SM was demonstrated in a novel manner by simultaneous two color optogenetic imaging with green (GCaMP6f) and red (RCaMP1.07) Ca^2+^ sensors expressed in ICC and SMCs, respectively. Imaging in this manner revealed the sequence of activation in ICC-SM and SMCs, showing clearly the frequency, onset and duration of Ca^2+^ transients in ICC-SM, the spatial spread of Ca^2+^ transients in ICC-SM networks, the development of Ca^2+^ transients in SMCs and tissue displacement (i.e. an optical indicator of muscle contraction). Correlation analysis demonstrated the coherence of these events. Ca^2+^ transients, lasting for about 2 sec propagated without decrement through networks of ICC-SM and preceded and likely initiated Ca^2+^ signaling and contractions in SMC, as was also suggested by intracellular microelectrode recordings from cells along the innermost surface of canine colonic muscles [40].

Slow waves with characteristics similar to those found in the stomach and small intestine (i.e. relatively fast upstroke depolarization and a plateau phase) are generated along the submucosal surface of the CM layer in the colon [12]. It was discovered that peeling the submucosa from the innermost surface of CM blocked slow waves [33]. While the authors of that study thought this tissue was mostly connective tissue with possibly some adherent SMCs, it is now clear that a population of pacemaker cells, ICC-SM, are present along the submucosal surface. ICC-SM and the networks they form are preserved and remain functionally similar in isolated submucosal tissues to ICC-SM attached to the muscularis. It should be noted that ICC-SM were more adherent to CM in the distal colon, and it was more difficult to obtain ICC-SM/submucosal preparations from that region. Isolation of submucosal tissues with adherent ICC-SM was used in the current study to eliminate movement artifacts generated by muscle contractions that plague high-resolution Ca^2+^ imaging in most smooth muscle tissues.

While the frequency of pacemaker activity was relatively stable over time in a given preparation, the sequence of activation of individual ICC-SM within the network varied as a function of time, as previously observed in gastric [86] and small intestinal [87] ICC-MY networks. What appeared as global Ca^2+^ transients in low resolution imaging partitioned into clusters of localized Ca^2+^ transients when viewed with a 60x objective at 30 frames per sec or at higher acquisition speeds. Summation of the clustered events reproduced the frequency and duration of the Ca^2+^ waves observed at low resolution. Multiple firing sites, averaging ∼8 per cell, were identified. This pattern of clustered Ca^2+^ transients was also observed in ICC-MY of the small intestine, the pacemaker cells in that region [88]. Organization of Ca^2+^ transients into clusters in ICC-SM was dependent upon voltage-dependent Ca^2+^ entry, and our data revealed that in contrast to ICC-MY of the small intestine, Ca^2+^ entry by both dihydropyridine-sensitive and insensitive mechanisms contributes to clustering and propagation of Ca^2+^ waves in intact networks. Nicardipine and isradipine reduced the occurrence of CTCs dramatically, and the T-channel antagonists, NNC 55-0396, TTA-A2 and Z-944 also reduced the occurrence and disordered the Ca^2+^ transients. *Cacna1c*, *Cacna1d* and *Cacna1h* were expressed in purified ICC-SM, and the presence and function of these channels can explain the pharmacological observations. Channels resulting from *Cacna1d* (encoding Ca_V_α1D) activate at relatively hyperpolarized membrane potentials and their currents are partially inhibited by dihydropyridines (∼ 50-70% of current density block) in comparison to *Cacna1c* gene products (Ca_V_α1C) [64, 89], but isradipine blocks Ca_V_α1C and Ca_V_α1D equally [90–92]. The fact that isradipine had no greater effect on the occurrence of clustered Ca^2+^ transients than nicardipine suggests that the L-type component of Ca^2+^ entry may be carried primarily by Ca_V_α1C channels. We have observed relatively robust expression of *Cacna1d* in a variety of ICC in mice [88, 93], and these channels may have a greater functional role at more negative resting membrane potentials.

Having three independent voltage-dependent Ca^2+^ conductances with different properties of voltage-dependent activation and inactivation coordinate clustering of Ca^2+^ transients provide a safety factor for preservation of pacemaker activity over a broad range of membrane potentials. In spite of overarching changes in membrane potential that might influence the availability of ion channels with narrow voltage-ranges, the broader range of activation potentials offered by expression and function of both L-type and T-type Ca^2+^ channels might protect against voltage-dependent inhibition of pacemaker activity. L-type channels are activated at less polarized potentials than T-type channels [94]. Thus, a factor producing tonic hyperpolarization of the SIP syncytium (e.g. purinergic inhibitory neurotransmission; [95, 96]) may tend to switch the dominant voltage-dependent Ca^2+^ entry mechanism from L-type to T-type Ca^2+^ channels. This concept was demonstrated by the decreased inhibitory effects of nicardipine and increased effects of NNC 55-0396 on Ca^2+^ transients after exposure of tissues to pinacidil. The opposite might be true if the SIP syncytium experiences a depolarizing influence (e.g. neurogenic or humerogenic).

Pinacidil hyperpolarizes colonic muscles through activation of K_ATP_ channels in SMCs [62]. This compound increased the frequency and decreased the duration of CTCs. These results are consistent with the effects of pinacidil on electrical pacemaker activity in the small intestine where it increases the d*V*/d*t*_max_ of the upstroke depolarization and decreases the durations of slow waves [56]. The increase in frequency may have been due to reduced inactivation and increased availability of Ca_V_α1D and T-type channels (Ca_V_α1H) at more hyperpolarized potentials. The decrease of duration of CTCs may be due to a relative shift in the importance of T-type vs. L-type channels with hyperpolarization. In the presence of pinacidil, NNC 55-0396 had increased antagonistic effects on CTCs. Ca^2+^ currents via T-type channels inactivate rapidly, whereas L-type channel inactivation is slower and incomplete [97–99]. Thus, the Ca^2+^ entry period for T-type currents is likely to be more transient than with L-type currents. Channel density in proximity to Ca^2+^ release channels may also affect the degree of coupling between Ca^2+^ entry and CICR, and as yet little is known about the structure and functional components of microdomains in ICC.

The importance of Ca^2+^ entry as the primary means of activation and organization of pacemaker activity in ICC-SM was shown by the dyscoordination of Ca^2+^ transients when extracellular Ca^2+^ was decreased and the incomplete effects of thapsigargin and CPA on Ca^2+^ transients. We noted tight clustering of Ca^2+^ transients at 2.5 and 2.0 mM [Ca^2+^]_o_ but the tightness of the CTCs disassociated when the driving force for Ca^2+^ entry (i.e. [Ca^2+^]_o_ was reduced to 1 mM), and frequency of CTCs was greatly reduced at concentrations lower than 1 mM. Our concept is that Ca^2+^ entry couples to CICR in ICC. Reducing the driving force for Ca^2+^ entry would be expected to reduce the probability for effective coupling to CICR. CICR would tend toward negligible when Ca^2+^ entry falls below threshold levels. Concentrations of thapsigargin and CPA that blocked Ca^2+^ transients quantitatively in other ICC [51, 81, 88] caused partial block of Ca^2+^ transients in ICC-SM. In fact, these drugs caused a marked narrowing of the duration of the CTCs, and this led us to analyze the temporal characteristics of Ca^2+^ transients within clusters. Ca^2+^ transients at the beginning of the CTCs were unaffected by ryanodine and xestospongin C, but transients toward the end of the clusters were blocked. These data suggest that the initial Ca^2+^ transients may result primarily from Ca^2+^ entry, and clusters are sustained temporally by Ca^2+^ release.

We have searched for a preparation of pacemaker ICC that would allow us to investigate the underlying pacemaker activity. We have speculated that stochastic Ca^2+^ release events, as occur in all ICC [15], are responsible for the spontaneous transient depolarizations (STDs) observed in patch clamp recordings from isolated ICC [16, 100]. No simultaneous recordings of Ca^2+^ transients and membrane currents or potentials changes have been achieved yet, and the expected link between Ca^2+^ transients and STDs is based on the fact that these events have common pharmacology and sensitivity to drugs that interfere with Ca^2+^ release [66]. Ca^2+^ transients and the spontaneous transient inward currents (STICs) due to activation of Ano1 channels result in spontaneous transient membrane depolarizations (STDs). Temporal summation of STDs is likely to be the generator potentials that activate T-type or L-type Ca^2+^ currents and initiate propagating slow wave events. In this concept it is logical to suggest that inhibition of Ca^2+^ release should reduce the duration of the CTCs, and inhibition of Ca^2+^ entry should inhibit the organizing influence of Ca^2+^ entry and block CTCs. When CTCs are blocked, stochastic Ca^2+^ transients may be unleashed, as occur in ICC-IM [81] and ICC-DMP [101] that lack expression of voltage-dependent Ca^2+^ entry mechanisms. Block of CTCs and unmasking of stochastic Ca^2+^ transients was accomplished by hyperpolarization with pinacidil and reduction in the availability of L-type and T-type Ca^2+^ channels with nicardipine and NNC 55-0396. ICC-SM, imaged as in this series of experiments provides a potent model for investigating basic pacemaker mechanisms and what happens to these events in response to neurotransmission, hormonal and paracrine inputs and pathological or inflammatory conditions.

Previous studies have supported a role for store-operated Ca^2+^ entry (SOCE) in maintaining Ca^2+^ release events in ICC [49, 81]. This is logical because Ca^2+^ release is extremely dynamic in ICC, and it is likely that store Ca^2+^ would be depleted without an effective recovery mechanism. SOCE depends upon the expression of Orai channels and the ER delimited activator of Orai, STIM, that senses ER Ca^2+^ and binds to and activates Orai channels when Ca^2+^ depletion of the ER occurs [102]. STIM and Orai are expressed in colonic ICC [103]. However, an antagonist of Orai, GSK7975A, reduced the frequency of CTCs, but failed to block these events at a concentration effective in blocking Ca^2+^ transients in small intestinal ICC-MY [49]. STIM/Orai interactions appear to have a role in Ca^2+^ store maintenance in ICC-SM, but our data suggest that Ca^2+^ entry via L-type and T-type Ca^2+^ channels also provide Ca^2+^ entry mechanisms that may contribute to store refilling. It could also be suggested that GSK7975A, a well-known antagonist for Orai1 (IC_50_ = 4 μM; [104]), is less effective on Orai2, the dominant isoform expressed in ICC-SM. However, studies on cortical neurons that express only Orai2 showed about 50% block of SOCE by GSK7975A (5 μM) [105].

In summary ICC-SM, as suggested from dissection and electrophysiological experiments [12], are pacemaker cells distributed in an electrically coupled network along the submucosal surface of the CM layer. Our experiments demonstrate that Ca^2+^ transients in ICC-SM couple to activation of Ca^2+^ transients and contractions in neighboring SMCs. The contractile events, called ‘ripples’ by some authors in describing integrated colonic contractions [106, 107], summate with the larger amplitude contractions emanating from the myenteric region of the *tunica muscularis* to produce mixing and propagated movements characteristic of colonic motility [108]. Data from this study suggest that voltage-dependent Ca^2+^ entry serves at least four important functions in the pacemaker activity of ICC-SM: i) Propagation of activity within the ICC-SM network depends upon voltage-dependent Ca^2+^ entry, and the functions and voltage-dependent properties of three types of Ca^2+^ conductances appear to provide a safety factor that tends to preserve pacemaker activity over a broad range of membrane potentials (see Fig. 17). ii) Ca^2+^ entry is the mechanism that organizes Ca^2+^ release events into CTCs. These events constitute the Ca^2+^ waves that propagate through ICC-SM networks, and presumably by activation of Ano1 channels cause slow wave depolarizations. iii) Ca^2+^ entry also appears to contribute to refilling of stores, as pacemaker activity was not as immediately dependent upon SOCE as in other ICC [49, 81]. iv) The observations that treatments expected to reduce Ca^2+^ release from stores and reduce coupling between Ca^2+^ entry and CICR reduced, but did not block CTCs, may indicate that transient Ca^2+^ entry (sparklets), possibly through activation of Ano1 and depolarization, may underlie the pacemaker functions of ICC-SM. Additional studies will be necessary to resolve these hypotheses in finer detail. The preparation of excised submucosal tissue with adherent ICC-SM removes movement artifacts from imaging and is likely to provide a powerful tool for improving resolution of pacemaker mechanisms and determining how regulatory and pathophysiological factors modify basic pacemaker mechanisms.

**Figure 17.**
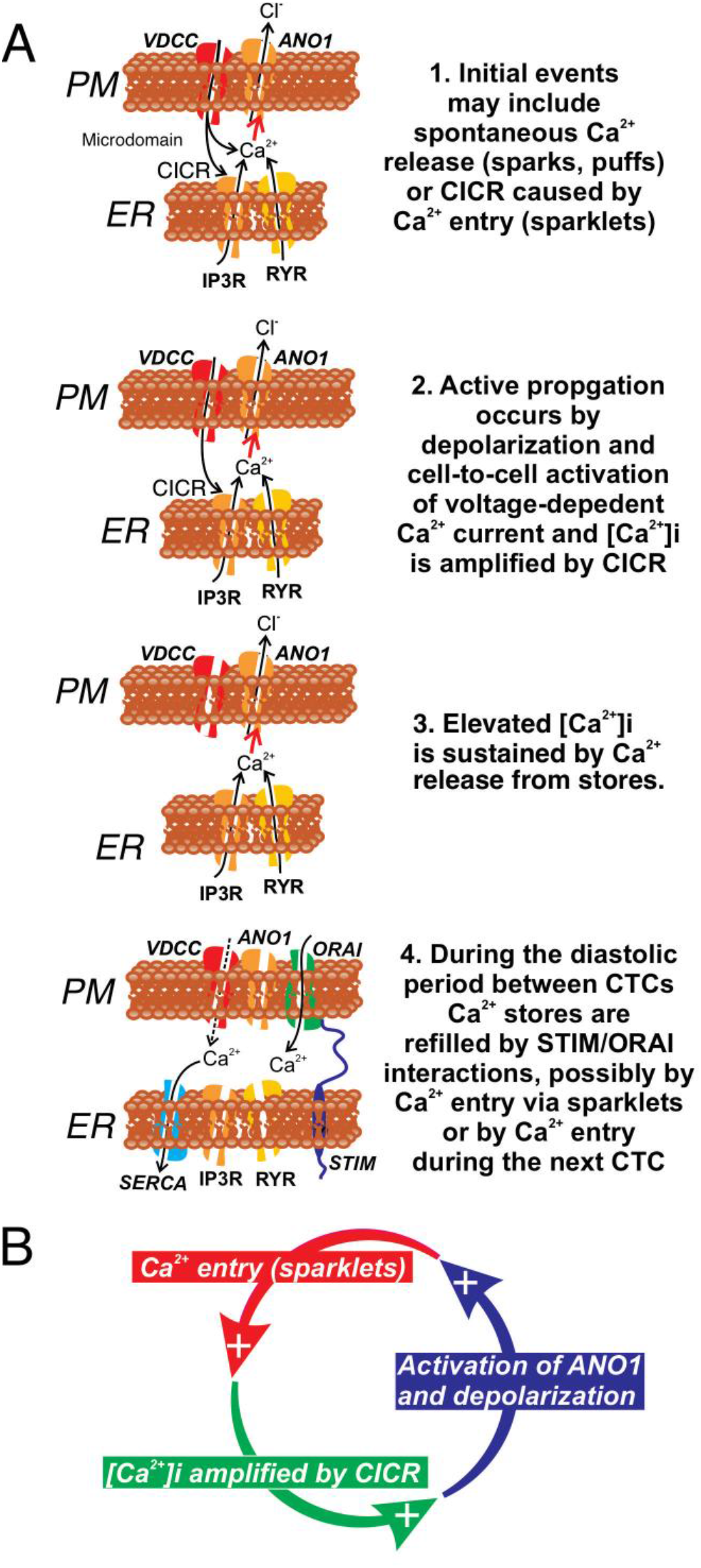
Role of voltage-dependent Ca^2+^ entry in the pacemaker function of ICC-SM. ***A*** Shows segments of plasma membrane (PM) and endoplasmic reticulum membrane (ER) that form PM-ER junctions and microdomains. At least 3 types of voltage-dependent Ca^2+^ channels (VDCC) are expressed in ICC-SM, Ca_V_1.2, Ca_V_1.3 and Ca_V_3.2. These conductances, with voltage-dependent activation and inactivation properties spanning a broad range of negative potentials, insure maintenance of pacemaker activity under conditions of hyperpolarization or depolarization in ICC-SM. Pacemaker activity (1. Initial events) in ICC-SM could be due to spontaneous release of Ca^2+^ from stores in the ER and utilize either IP_3_R or RYR receptors or both (Ca^2+^ sparks and puffs). However, our data cannot rule out the possibility that transient openings of voltage-dependent Ca^2+^ channels (sparklets) and amplification of Ca^2+^ in microdomains by CICR constitute the initial events of pacemaker activity. In this case Ca^2+^ release from stores is not the primary pacemaker event but a secondary response to Ca^2+^ entry. Inhibition of Ca^2+^ release from stores would lead to reduced probability of CICR and decrease the frequency of CTCs. Our hypothesis is that Ca^2+^ entry and/or release from stores activates Ca^2+^-dependent Cl^-^ current due to ANO1 in the plasma membrane. Active propagation between cells in ICC networks (Phase 2) was inhibited by blocking voltage-dependent Ca^2+^ channels. Active propagation may also require or depend upon amplification of Ca^2+^ in microdomains by CICR. The duration of Ca^2+^ entry is likely to be brief due to voltage-dependent inactivation of L-and T-type Ca^2+^ channels. The duration of CTCs appears to be enhanced by CICR (Phase 3). Our data show that the duration of CTCs is reduced by several manipulations known to inhibit Ca^2+^ release from stores. In Phase 4 store reloading may occur by multiple mechanisms and may include: i) transient Ca^2+^ entry via sparklets, ii) activation of SOCE via STIM/ORAI interactions, and iii) the increase in Ca^2+^ entry that occurs via depolarization and activation of Ca^2+^ entry at the onset of each CTC. ***B*** A novel hypothesis emerges from this study suggesting that the pacemaker mechanism in non-voltage-clamped cells includes a cyclical, positive-feedback phenomenon that may be responsible for initiation of CTCs and relies on: i) Ca^2+^ entry through voltage-dependent Ca^2+^ channels. Openings of clusters of these channels would generate sparklets; ii) Ca^2+^ entry initiates CICR which amplifies [Ca^2+^]_i_ within microdomains; iii) the rise in [Ca^2+^]_i_ activates ANO1 channels in the PM causing depolarization; iv) depolarization enhances the open probability of voltage-dependent Ca^2+^ channels, increasing Ca^2+^ entry. This cycle creates positive feedback for Ca^2+^ entry, clustering of localized Ca^2+^ transients due to Ca^2+^ entry during the first 350-450 ms of CTCs and development of slow wave depolarizations in ICC-SM.

## Methods

### Animals

Kit*^+/copGFP^* mice (B6.129S7-*^Kittm1Rosay/J^*; 5-8 wk old) were bred in house (Ro et al. 2010). GCaMP6f-floxed mice (Ai95 (RCL-GCaMP6f)-D) and C57BL/6 mice, their wild-type siblings, were purchased from Jackson Laboratories (Bar Harbor, MN, USA). Kit-iCre mice (c-Kit^+*/Cre-ERT2*^) were gifted from Dr. Dieter Saur (Technical University Munich, Munich, Germany).

#### Generation of Kit-iCre-GCaMP6f/Acta2-RCaMP1.07 mice

*Acta2-RCaMP1.07* mice (*tg(RP23-370F21-RCaMP1.07)B3-3Mik/J*) express the fluorescent Ca^2+^ indicator RCaMP1.07 in smooth muscle cells under the control of the Acta2 locus promoter/enhancer regions were obtained from Jackson Laboratories (Bar Harbor, MN, USA). To generate cell-specific expression in 2 distinct cell types (ICC and smooth muscle cells) *Acta2-RCaMP1.07* mice were bred with *Kit^Cre-ERT2^/GCaMP6f^fl/fl^* mice. The offspring *Kit-iCre-GCaMP6f/Acta2-RCaMP1.07* mice were identified by genotyping after receiving tamoxifen which served to delete the STOP cassette in the Cre-expressing cells; resulting in the expression of the fluorescent Ca^2+^ indicator protein, GCaMP6f. These mice allowed simultaneous, dual color imaging of ICC and smooth muscle cells.

iCre mice were injected with tamoxifen (TAM; Intraperitoneal injection; IP) at 6-8 weeks of age (2 mg of TAM for three consecutive days), as described previously (Baker *et al*., 2016), to induce activation of the Cre recombinase and expression of optogenetic sensors. Mice were used for experiments 10-15 days after the tamoxifen injections. On days of experiments the mice were anaesthetized by inhalation of isoflurane (Baxter, Deerfield, IL, USA) and killed by cervical dislocation before excision of tissues. The animals used, protocols performed and procedures in this study were in accordance with the National Institutes of Health Guide for the Care and Use of Laboratory Animals and approved by the Institutional Animal Use and Care Committee at the University of Nevada, Reno.

### Tissue preparation

Colonic segments (2 cm in length, proximal region) were removed from mice after an abdominal incision and placed in Krebs-Ringer bicarbonate solution (KRB). The tissues were cut along the mesenteric border and intraluminal contents were washed away with KRB. Tissues were prepared by blunt dissection in two ways: 1) The submucosa layer was isolated after carefully removing the mucosal layer and the *tunica muscularis* 2) The submucosa layer was left attached to the *tunica muscularis* after removal of the mucosa. The isolated submucosal layer preparation provided better imaging of ICC-SM by eliminating motion artifacts associated with muscle contractions. We used the isolated submucosal layer preparation in most cases in this study with the exception of two experiments where muscle attachments were necessary to test important questions (see Results).

### Immunohistochemistry

Colonic tissues from wild-type mice were processed to assess distribution of c-Kit immunoreactivity. Whole mounts of submucosal layer after removing the mucosa and *tunica muscularis* were fixed in 4% paraformaldehyde and visualized as described previously [22]. Briefly, after block with 1% bovine serum albumin, colonic tissues were incubated with a polyclonal antibody raised against c-Kit (mSCFR, R&D Systems, MN, USA; 1:500 dilution in 0.5% Triton-X working solution) for 48 h. Immunoreactivity was detected using Alexa-488 labeled donkey anti-goat IgG (1:1000 in PBS; Invitrogen, NY, USA). Colonic tissues were visualized using a Zeiss LSM 510 confocal microscope and images were constructed using Image J software (National Institutes of Health, MD, USA, http://rsbweb.nih.gov/ij). ICC (copKit) images were visualized using spinning-disk confocal system (CSU-W1; spinning disk, Yokogawa Electric, Tokyo, Japan).

### Cell sorting and quantitative PCR

Kit^+*/copGFP*^ mice (B6.129S7-*^Kittm1Rosay/J^*; 5-8 wks old) were used for evaluations of gene expression in ICC-SM. Cell-specific expression of the fluorescent reporter allows unequivocal identification of ICC [43]. After cell dispersion, ICC-SM were sorted by fluorescence-activated cell sorting (FACS) and evaluated for purity as previously described (Baker *et al*., 2016). Total RNA was isolated using an Illustra RNAspin Mini RNA Isolation Kit (GE Healthcare). qScript cDNA SuperMix (Quanta Biosciences), used according to the manufacturer’s instructions, was used to synthesize first-strand cDNA. Quantitative PCR (qPCR) was performed using Fast Sybr Green chemistry on the 7900HT Fast Real-Time PCR System (Applied Biosystems) and gene-specific primers (Supplemental Table 1). Regression analysis was performed to generate standard curves from the mean values of technical triplicate qPCRs of log10 diluted cDNA samples. Evaluation of gene expression in ICC-SM was compared with expression in the unsorted cells from the submucosal tissue of Kit^+*/copGFP*^ mice

### Ca^2+^ imaging

The isolated/intact submucosal layers were pinned to Sylgard coated dish and perfused with KRB solution at 37°C for a 60 min equilibration period. Ca^2+^ imaging was performed using a spinning-disk confocal system (CSU-W1; spinning disk, Yokogawa Electric, Tokyo, Japan) mounted on an upright Nikon Eclipse FN1 microscope equipped with several water immersion Nikon CFI Fluor lenses (10× 0.3 NA, 20× 0.5 NA, 40× 0.8 NA, 60× 0.8 NA and 100× 1.1 NA) (Nikon Instruments, New York, USA). The system is equipped with two solid-state laser lines of 488 nm and 561 nm. The laser lines are combined with a borealis system (ANDOR Technology, Belfast, UK) to increase laser intensity and uniformity throughout the imaging field of view (FOV). The system also has two high-speed electron multiplying charged coupled devices (EMCCD) cameras (Andor iXon-Ultra 897 EMCCD Cameras; ANDOR Technology, Belfast, UK) to allow dual-color imaging simultaneously and maintain sensitive and fast speed acquisition at full frame of 512×512 active pixels as previously described (Baker *et al*., 2015). Briefly, images were captured, and image sequences were collected at 33 to 50 fps using MetaMorph software (MetaMorph Inc., TN, USA). Experiments with pharmacological agents, a control activity period of (30 sec) was recorded prior of drug application into the chamber for 15 minutes.

### Ca^2+^ imaging analysis

Movies of Ca^2+^ transients in ICC-SM were imported, preprocessed and analyzed using a combination of 3 image analysis programs: 1) custom build software (Volumetry G8d, Dr. Grant Hennig); 2) Fiji/Image J (National Institutes of Health, MD, USA, http://rsbweb.nih.gov/ij) 3) Automated Spatio Temporal Map analysis plugin (STMapAuto), https://github.com/gdelvalle99/STMapAuto as described previously [44–46]. Briefly, movies of Ca^2+^ transients (stacks of TIFF images) were imported into Volumetry G8d and motion stabilized, background subtracted and smoothed (Gaussian filter: 1.5 x 1.5µm, StdDev 1.0). A particle analysis routine was employed using a flood-fill algorithm to enhance Ca^2+^ transient detection. particles (PTCLs) representing the areas of active Ca^2+^ signals in cells were saved as a coordinate-based particle movie and combined area and total number of PTCLs were calculated. To better isolate firing sites, only those particles that did not overlap with any particles in the previous frame but overlap with particles in the subsequent 70 ms were considered firing/initiation sites.

### Drugs and solutions

All tissues were perfused continuously with KRB solution containing (mmol/L): NaCl, 5.9; NaHCO_3_, 120.35; KCl, 1.2; MgCl_2_, 15.5; NaH_2_PO_4,_1.2; CaCl_2_, 2.5; and glucose, 11.5. The KRB solution was warmed to a physiological temperature of 37 ± 0.3 °C and bubbled with a mixture of 97 % O_2_ – 3 % CO_2_. For experiments utilizing external solutions with 0 [Ca^2+^]_o_, CaCl_2_ was omitted and 0.5 mM ethylene glycol-bis (β-aminoethyl ether)-N, N, N’, N’– tetraacetic acid (EGTA) was added to the solution. NNC 55-0396 and TTA-A2 were purchased from Alomone Labs (Jerusalem, Israel). 2-aminoethyl-diphenylborinate (2-APB), tetracaine, nicardipine, pinacidil was purchased from Millipore-Sigma (St. Louis, Missouri, USA). Thapsigargin, isradipine, Z-944, CPA and ryanodine were purchased from Tocris Bioscience (Ellisville, Missouri, USA). GSK 7975A was purchased from Aobious (Aobious INC, MA, USA), and xestospongin C (XeC) was purchased from Cayman Chemical (Michigan, USA).

### Statistical analysis

Data is presented as the mean ± standard error unless otherwise stated. Statistical analysis was performed using either a Students *t*-test or one-way ANOVA with a Tukey post hoc test where appropriate. In all tests, P<0.05 was considered significant. When describing data, *n* refers to the number of animals used in a dataset. Probabilities < 0.05 are represented by a single asterisk (*), probabilities < 0.01 are represented by two asterisks (**), probabilities < 0.001 are represented by three asterisks (***) and probabilities < 0.0001 are represented by four asterisks (****). All statistical tests were performed using GraphPad Prism 8.0.1 (San Diego, CA).

## Abbreviations

CM: Circular Muscle
FOV: Field of view
GI: Gastrointestinal
ICC: Interstitial Cells of Cajal
ICC-DMP: Interstitial cells of Cajal at the level of the deep muscular plexus
ICC-IM: Intramuscular interstitial cells of Cajal
ICC-MY: Interstitial cells of Cajal at the level of the myenteric plexus
ICC-SM: Interstitial cells of Cajal at the submucosal border
GCaMP: Genetically encoded Ca^2+^ indicator composed of a single GFP
IP_3_: Inositol 1,4,5-trisphosphate
InsP_3_R: Inositol triphosphate receptor
KRB: Krebs Ringer Bicarbonate
LM: Longitudinal Muscle
PDGFRα: Platelet derived growth factor receptor α
ROI: Region of interest
RyR: Ryanodine receptor
SIP: syncytium Electrical syncytium formed by **S**mooth muscle cells, **I**CC and **P**DGFRα^+^ cells in GI muscles
SERCA: Sarco/endoplasmic reticulum Ca^2+^-ATPase
SMC: Smooth muscle cell

## Additional information

### Author contributions

1. Conception and design of the study: SAB, KMS
2. Collection and analysis of data: SAB, WAL, IFD, CAC, BTD, KMS
3. Interpretation of data: SAB, BTD, KMS
4. Drafting the article: SAB, KMS
5. Revising article critically for intellectual content: SAB, BTD, CAC, SMW, KMS All authors read and approved the manuscript before submission. All persons designated as authors qualify for authorship, and all those who qualify for authorship are listed. All authors agree to be accountable for all aspects of the work in ensuring that questions related to the accuracy or integrity of any part of the work are appropriately investigated and resolved

## Acknowledgements

The authors would like to extend their sincere appreciation to Yulia Bayguinov for assistance with immunohistochemical experiments, Lauren O’Kane for assistance with qPCR experiments and David White and Emily Fox for assistance with FACS.

## Supplemental material

**Supplemental Movie 1: Simultaneous dual color imaging of ICC-SM and SMCs in the colon**

A video of propagating Ca^2+^ waves through an ICC-SM network in proximal colon of *Kit-iCre-GCaMP6f/Acta2-RCaMP1.07* strain imaged with a 20× objective. simultaneous imaging of 2 optogenetic Ca^2+^ sensors: *GCaMP6f* in ICC-SM (left FOV; green) and *RCaMP1.07* in SMCs (right FOV; red) with different fluorescence characteristics (ensuring minimal spectral overlap). the signal coordination between ICC and SMCs showing the correlation between Ca^2+^ transients in the ICC-SM network and activation of SMCs adjacent to ICC-SM. ICC-SM transients (green trace) preceded Ca^2+^ signals in SMCs (red trace). The scale bar (yellow) is 25 µm.

**Supplemental Movie 2: High spatial resolution ICC-SM Ca2^+^ signals composed of multiple Ca2^+^ firing sites**

**A** video showing subcellular Ca^2+^ transients in ICC-SM at high resolution imaged with a 60× objective. Ca^2+^ signals were monitored using the genetically encoded Ca^2+^ indicator GCaMP6f. The left panel shows typical stellate-shaped ICC-SM with multiple interconnected processes. The scale bar (yellow) is 10 µm. The middle panel shows the Ca^2+^ particle (PTCL) activity, color coded in blue for raw PTCLs, and the centroids of particles are indicated in purple and in green indicate Ca^2+^ firing sites. Note the multiple-site firing of Ca^2+^ transients in ICC-SM. The right panel shows initiation/firing sites accumulation map. The pattern of firing sites Ca^2+^ activity was temporally clustered as Ca^2+^ wave oscillations sweeps through ICC-SM networks.

---

**Supplemental Figure 1.**
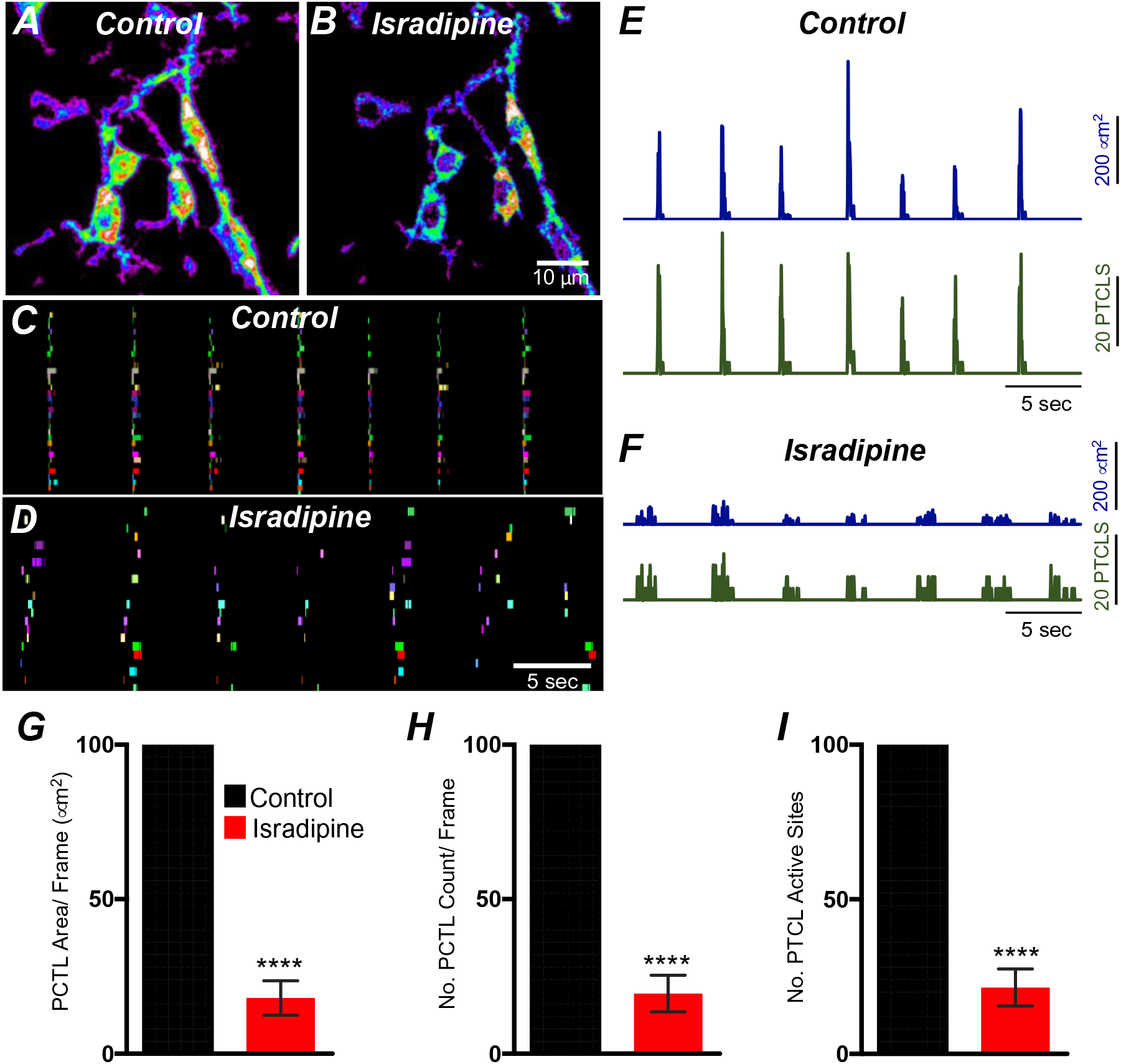
Isradipine effects on Ca^2+^ transients in ICC-SM. ***A&B*** Heat-map images showing the summated Ca^2+^ transient particles of ICC-SM under control conditions and in the presence of isradipine (1μM). ***C&D*** Occurrence maps of individually color-coded Ca^2+^ firing sites in the ICC-SM in the FOV under control conditions and in isradipine. Ca^2+^ transient PTCL activity Traces over an entire recording of the ICC-MY showing PTCL area/frame (blue) and PTCL count/frame (green) in control conditions ***E*** and in the presence of isradipine (1μM) ***F***. Summary graphs of average percentage change of PTCL area ***G***, PTCL count ***H*** and the number of Ca^2+^ firing sites was significantly affected by isradipine ***I*** (*n*= 7). Significance determined using unpaired t-test, **** = *P*<0.0001. All data graphed as mean ± SEM.

**Supplemental Figure 2.**
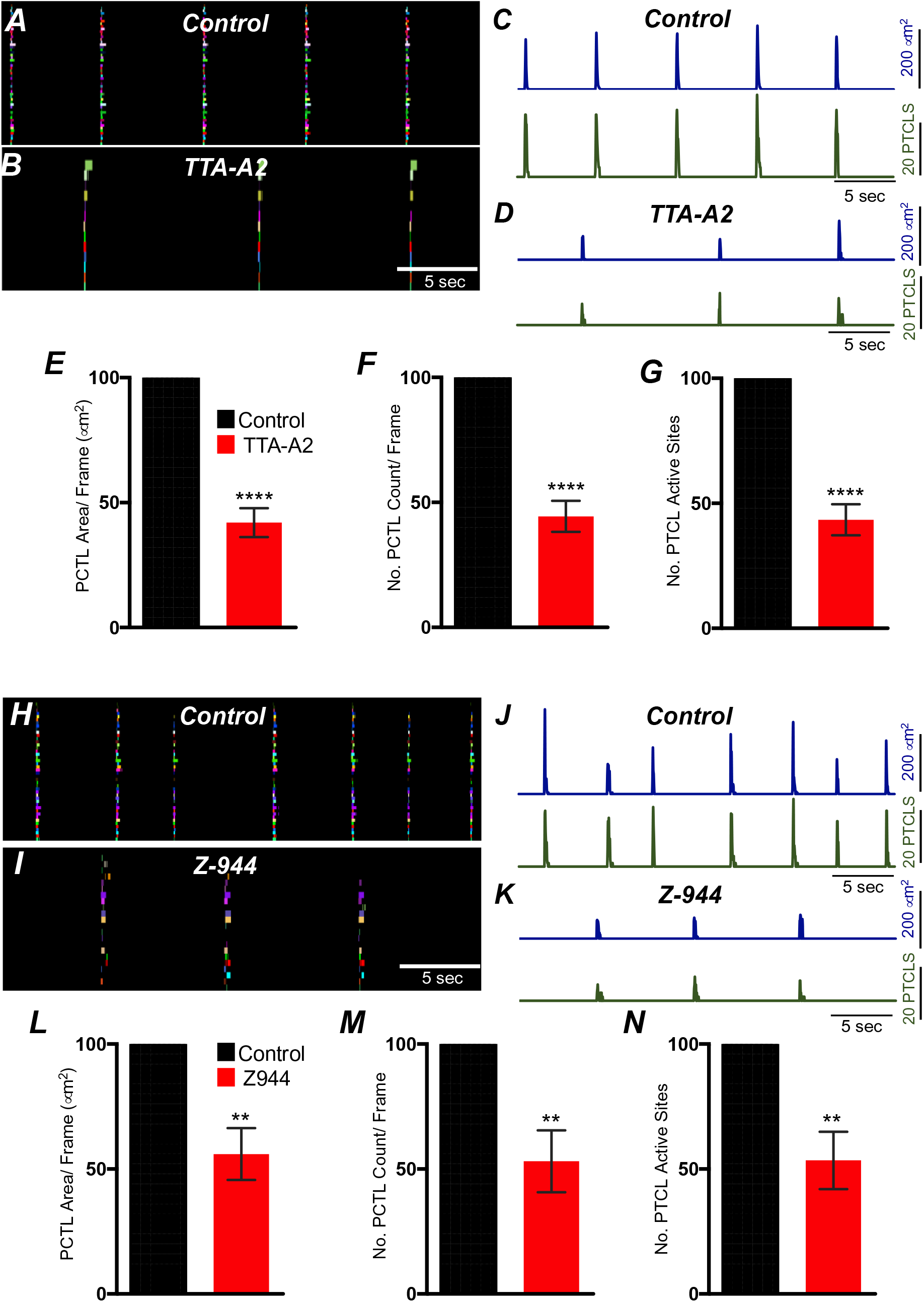
Effects of T-type Ca^2+^ channel antagonists, TTA-A2 and Z-944 on ICC-SM Ca^2+^ transients. ***A&B*** Occurrence maps showing ICC-SM network active firing sites. Sites were individually color-coded and plotted under control ***A*** and in the presence of TTA-A2 (10 μM) ***B*** conditions. Plots of Ca^2+^ transient particle activity of ICC-SM in control conditions and in the presence of TTA-A2 showing PTCL area (blue) and PTCL count (green) under control conditions ***C*** and in the presence of TTA-A2 ***D***. Summary graphs of average percentage change of PTCL area ***E***, PTCL count ***F*** and the number of Ca^2+^ firing sites was reduced by TTA-A2 ***G*** (*n*= 7). ***H&I*** Active firing sites were individually color-coded and plotted as an occurrence maps in the ICC-SM network under control ***H*** and Z-944 (1 μM) ***I*** conditions. Plots of Ca^2+^ transient particle activity of ICC-SM in control conditions and in the presence of Z-944 showing PTCL area (blue) and PTCL count (green) under control conditions ***J*** and in the presence of Z-944 ***K***. Summary graphs of average percentage change of PTCL area ***L***, PTCL count ***M*** and the number of Ca^2+^ firing sites was reduced by Z-944 ***N*** (*n*= 5). Significance determined using unpaired t-test, ** = *P*<0.01,**** = *P*<0.0001. All data graphed as mean ± SEM.

**Supplemental Figure 3.**
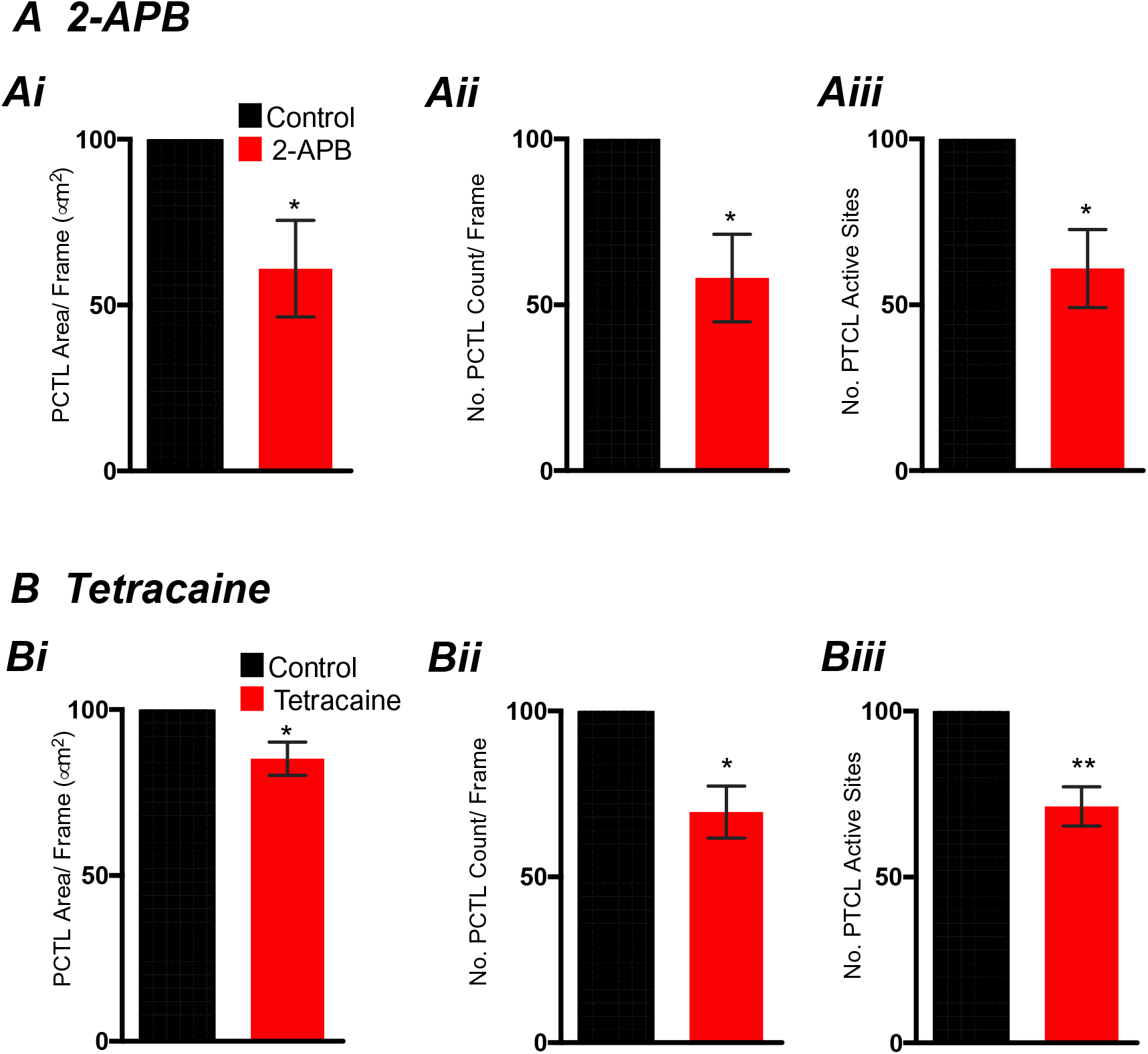
Effects of 2-APB and Tetracaine on ICC-SM Ca^2+^ transients. ***A*** Summary graphs of Ca^2+^ activity in ICC-SM in the presence of 2-APB (100 μM) are shown in ***Ai*** (PTCL area), ***Aii*** (PTCL count) and ***Aiii*** the number of PTCL active sites (*n*= 5). ***B*** Summary graphs of Ca^2+^ activity in ICC-SM in the presence of tetracaine (100 μM) are shown in ***Bi*** (PTCL area), ***Bii*** (PTCL count) and ***Biii*** the number of PTCL active sites (*n*= 5). Significance determined using unpaired t-test, * = *P*<0.1,** = *P*<0.01. All data graphed as mean ± SEM.

**Supplemental Table 1:**
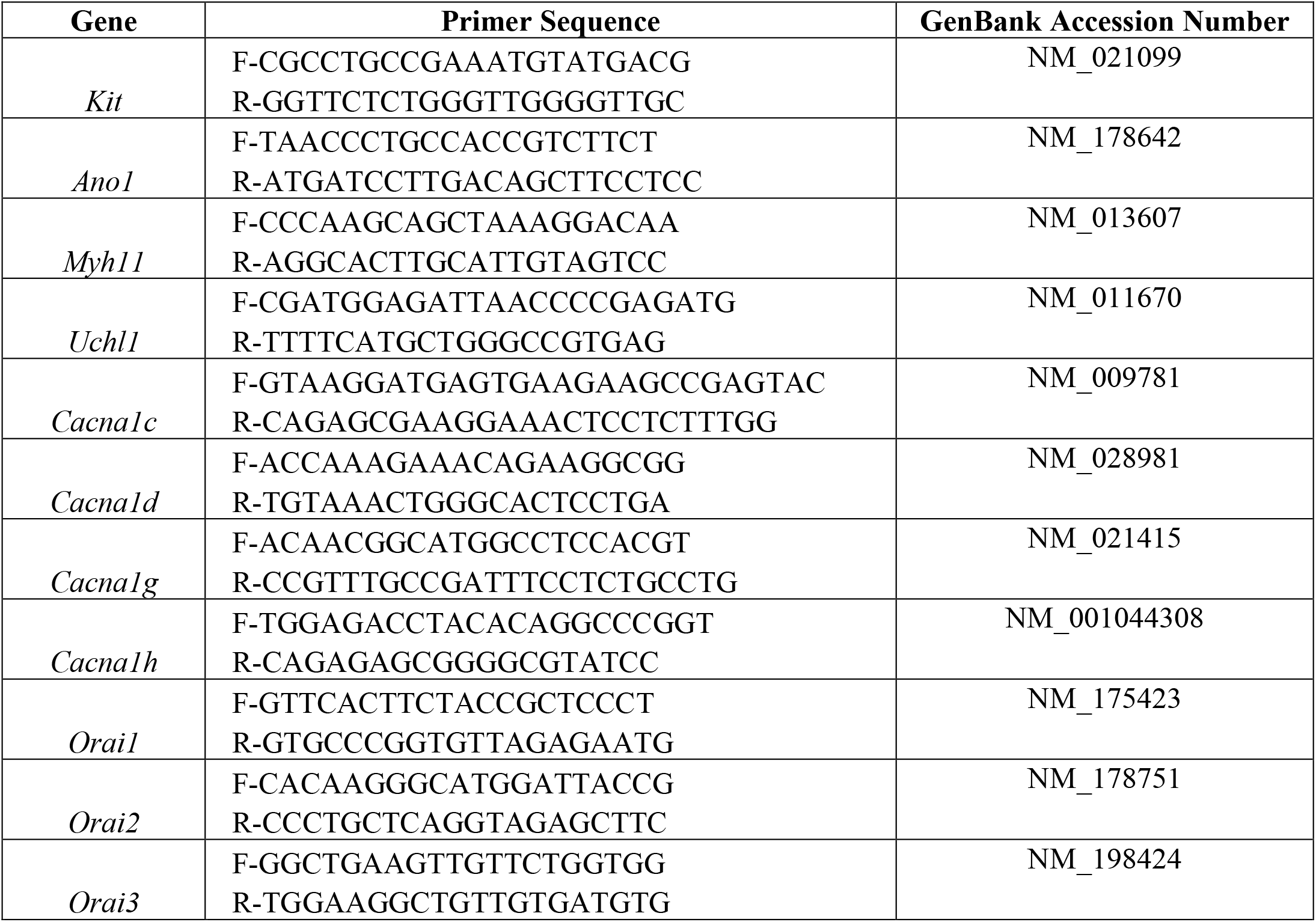
Summary of gene primers sequences of Kit, Ano1, Myh11, Uchl1, Cacna1c, Cacna1d, Cacna1g, Cacna1h and Orai1-3

